# The MRTFs are required for thymocyte trafficking and T_reg_-mediated immune tolerance

**DOI:** 10.64898/2026.10.02.756229

**Authors:** Patrick Costello, Diane Maurice, Maria Benito-Jardon, Richard Treisman

**Author notes:** Autoimmunity Laboratory, Francis Crick Institute, 1 Midland Road, London NW1 1AT, UK.

## Abstract

The MRTFs (myocardin-related transcription factors) are essential for effective cytoskeletal function during differentiation and development. To investigate the role played by the MRTFs in T cell development and function we inactivated *Mrtfb* in *Mrtfa*-null mice at the DN2 stage (*Mrtfab*KO^T^ animals). Positive selection, and development of *Mrtfab*KO^T^ regulatory (T_reg_), γδ, and NK T cells proceeded normally. Thymic egress was impaired, however, with mature T cells accumulating at the cortico-medullary junction, and *in vitro Mrtfab*KO^T^ thymocytes exhibited impaired cytoskeletal responses to S1P and migratory cytokines. *Mrtfab*KO^T^ mice were lymphopenic, with defective homeostatic proliferation and T cell homing to peripheral lymphoid organs. Although thymocyte negative selection was unaffected, *Mrtfab*KO^T^ animals develop autoimmune phenotypes in lacrimal gland and intestinal mucosa. Using the experimental colitis model, we show that these phenotypes reflect functional deficits in *Mrtfab*KO^T^ T_reg_ cells. Our findings show that MRTF-SRF signalling plays an unanticipated role in thymocyte egress and immune tolerance, and reveal a role of the MRTFs in T_reg_ function.

## INTRODUCTION

The production of self-tolerant functional T cells by the thymus involves multiple signalling events mediated by the TCR and chemokine receptors, which coordinate cell survival, proliferation, differentiation and migration (for reviews see Ashby and Hogquist, 2024). Following thymic entry at the cortico-medullary junction (CMJ), a process strongly dependent on CCR9, thymocytes differentiate as they migrate through the cortex. Developing thymοcytes then undergo a series of selection processes that generate functional and self-tolerant T cells and regulatory T cell (T_reg_) lineages (Ashby and Hogquist, 2024). Successful assembly of the αβTCR allows thymocyte positive selection and differentiation to CD4+ or CD8+ SP thymocytes.

Following positive selection, thymocytes migrate into the medulla. Here, signalling induced by high-affinity TCR-MHC-peptide interaction with stromal cells induces deletion of potentially self-reactive thymocytes, whereas thymocytes receiving relatively strong, but sub-threshold, TCR signals are preferentially diverted into the T_reg_ lineage (Gascoigne et al., 2016; Klein et al., 2014). Migration into and within the medulla is critically dependent on CCR7, whose ablation leads to defects in central tolerance (Forster et al., 2008; Worbs and Forster, 2007). After further maturation, thymocytes egress to the periphery by reverse-endothelial migration, again at the CMJ, a process dependent on S1PR signalling (Cyster and Schwab, 2012). Tolerance is ensured by two principal mechanisms: negative selection of high-affinity autoreactive TCRs, and peripheral suppression by T_regs_ (Ashby and Hogquist, 2024).

The SRF (Serum response factor) transcription factor network integrates proliferative and cytoskeletal signals, and is intimately involved in many developmental and disease settings that involve signal-regulated cytoskeletal dynamics. SRF activity is controlled by two families of signal-related co-factors: the TCFs (ternary complex factors; SAP-1, Elk-1, Net), controlled by Ras-ERK signalling, and the MRTFs (myocardin related transcription factors), which couple transcription to actin dynamics (for overview, see Olson and Nordheim, 2010; Posern and Treisman, 2006). In T cells, MRTF-SRF signalling is required for the sustained proliferative response of CD8+ T cells to L monocytogenes infection, facilitating homotypic cell interactions and IL-2 paracrine signalling (Maurice et al., 2024), while in hematopoietic stem cells it is required for bone marrow colonisation (Costello et al., 2015). The role of MRTF-SRF signalling in T cell development has not been investigated, but TCF-SRF signalling is necessary and sufficient for positive selection (Costello et al., 2010; Costello et al., 2004; Mylona et al., 2011).

Here we investigated the role of MRTF-SRF signalling in T cell development. We find that MRTF-null thymocytes undergo positive selection normally, but exhibited reduced cytoskeletal responses to stimuli acting through the SDF-1, CCL19, CCL21, and S1P receptors. Consistent with this, they exhibit defects in thymocyte progenitor homing and mature thymocyte egress, disrupted thymic architecture. These phenotypes are associated with profound lymphopenia and defects in central tolerance, and we show that the from T_reg_ functional deficits. The two arms of the SRF network thus play distinct roles in thymocyte differentiation and T cell function.

## RESULTS

### The MRTFs are not required for thymocyte positive selection

We previously showed that inactivation of all 3 TCF cofactors severely impairs but does not abolish thymocyte positive selection, although SRF is absolutely required, suggesting that MRTF-SRF signalling might contribute to thymocyte selection (Costello et al., 2010; Mylona et al., 2011). To investigate the role of the MTRFs detail we crossed *Mrtfa*^-/-^*Mrtfb*^f/f^ (Li et al., 2006; Mokalled et al., 2015) mice with mice expressing Cre recombinase from human CD2 minigene promoter (hCD2-Cre, de Boer et al., 2003). In the resulting *Mrtfa*^-/-^ *Mrtfb*^f/f^ *hCD2-Cre* offspring, thymocytes lack both *Mrtfa and Mrtfb* from the DN2 stage onwards (hereafter referred to as *Mrtfab*KO^T^ cells; Figure S1A). Thymic cellularity of *Mrtfab*KO^T^ animals was approximately 50% reduced compared to age- and sex-matched hCD2Cre controls (Figure S1B). All CD4-CD8-(DN) populations were represented in *Mrtfab*KO^T^ thymus, but the proportion and numbers of CD44+CD25-DN1 cells was greatly reduced (Figure 1A), as were the proportion and number of CD44+ CD25-CD117+ early thymocyte precursors (ETPs; Figure 1A). A comparable reduction in cellularity and ETP numbers was observed in *Mrtfa*^-/-^ thymus (Figure S1B, S1C), indicating that MRTF activity is limiting for effective ETP thymic homing.

**Figure 1.**
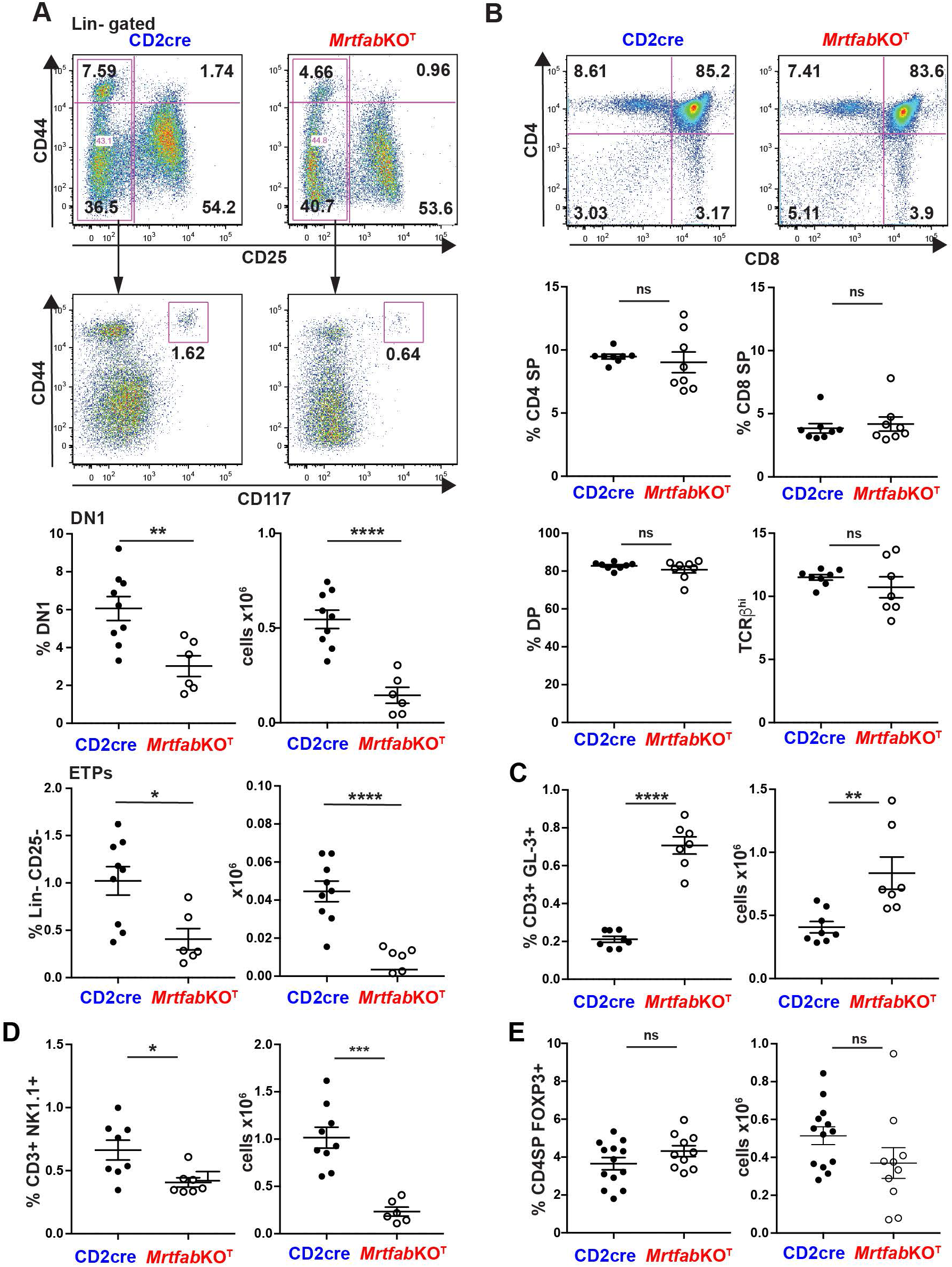
The MRTFs are required for thymic homing but not positive selection. A. Top, representative live-gated, lineage-negative CD44 CD25 profiles; middle, CD117 CD44 profiles of the CD25-CD44^lo^ populations, with CD44+ CD117+ ETPs indicated. Bottom, DN1 and ETP quantitation, combining two independent experiments, at least 3 female mice each genotype. B. Top, representative live-gated CD4 and CD8 profiles of CD2cre and *Mrtfab*KO^T^ thymocytes, with proportions of CD4+, CD8+, and DP shown. Bottom, quantitation of CD4SP, CD8SP, DP and TCRβhi, combining two independent experiments, at least 3 mice each genotype. C. Proportions and numbers of CD3+ GL3+ γδ thymocytes in CD2cre and *Mrtfab*KO^T^ mice. Data combined from two independent experiments, at least 3 mice each genotype. D. Proportions and numbers of CD3+ NK1.1+ thymic NK T cells in CD2cre and *Mrtfab*KO^T^ mice. Data combined from two independent experiments, at least 3 mice each genotype. E. Proportions and numbers of Foxp3+ CD4+ thymocytes in CD2cre and *Mrtfab*KO^T^ mice. Data combined from three independent experiments, at least 3 mice each genotype.

Although overall thymus cell numbers were decreased, the proportions of DP, CD4SP and CD8SP thymocytes, and TCRβ^hi^ cells in *Mrtfab*KO^T^ thymus were comparable to controls (Figure 1B), and this was also seen with TCR transgenes, as will be seen below. Similar results were seen in *Mrtfa*^-/-^thymus (Figure S1D). Strikingly, both the proportion and numbers of γδ thymocytes were increased ∼3-fold in *Mrtfab*KO^T^ thymus despite the reduced cellularity (Figure 1C), and a slight increase in γδ thymocytes was also seen in *Mrtfa*^-/-^ thymus (Figure S1E). The proportion and numbers of mature CD3+ NK1.1+ cells were significantly reduced in *Mrtfab*KO^T^ thymus (Figure 1D). Finally, the proportion of Foxp3+ CD4 SP T_reg_ thymocytes was slightly increased, and the numbers correspondingly decreased, although this was not statistically significant (Figure 1E, Figure S1F). Taken together, these data show that MRTF inactivation does not impair thymocyte positive selection.

### MRTF-SRF signalling is required for thymocyte thymic egress

In LFA1^-/-^ and S1PR^-/-^ animals, elevated γδ cell numbers are associated with defects S1P-dependent thymic egress, and are accompanied by increased numbers of thymocytes expressing maturation markers (Allende et al., 2004; Bose et al., 2014; Matloubian et al., 2004; Odumade et al., 2010). Consistent with this, total TCR^hi^ thymocytes contained increased proportions of Qa2^hi^ CD24^lo^ cells (**Figure S2A**). Analysis of positively-selected TCR^hi^ thymocytes with γδ, T_reg_ and NK1.1^+^ cells excluded revealed a substantially increased proportion of mature CD69-CD62L+ and M2 CD69-MHCI+ cells (**Figure 2A, S2B**). Similar results were obtained using TCR transgenes (see below). A similar but less pronounced increase in mature thymocytes was observed in MRTF-A null animals (**Figure S2C, S2D**).

**Figure 2.**
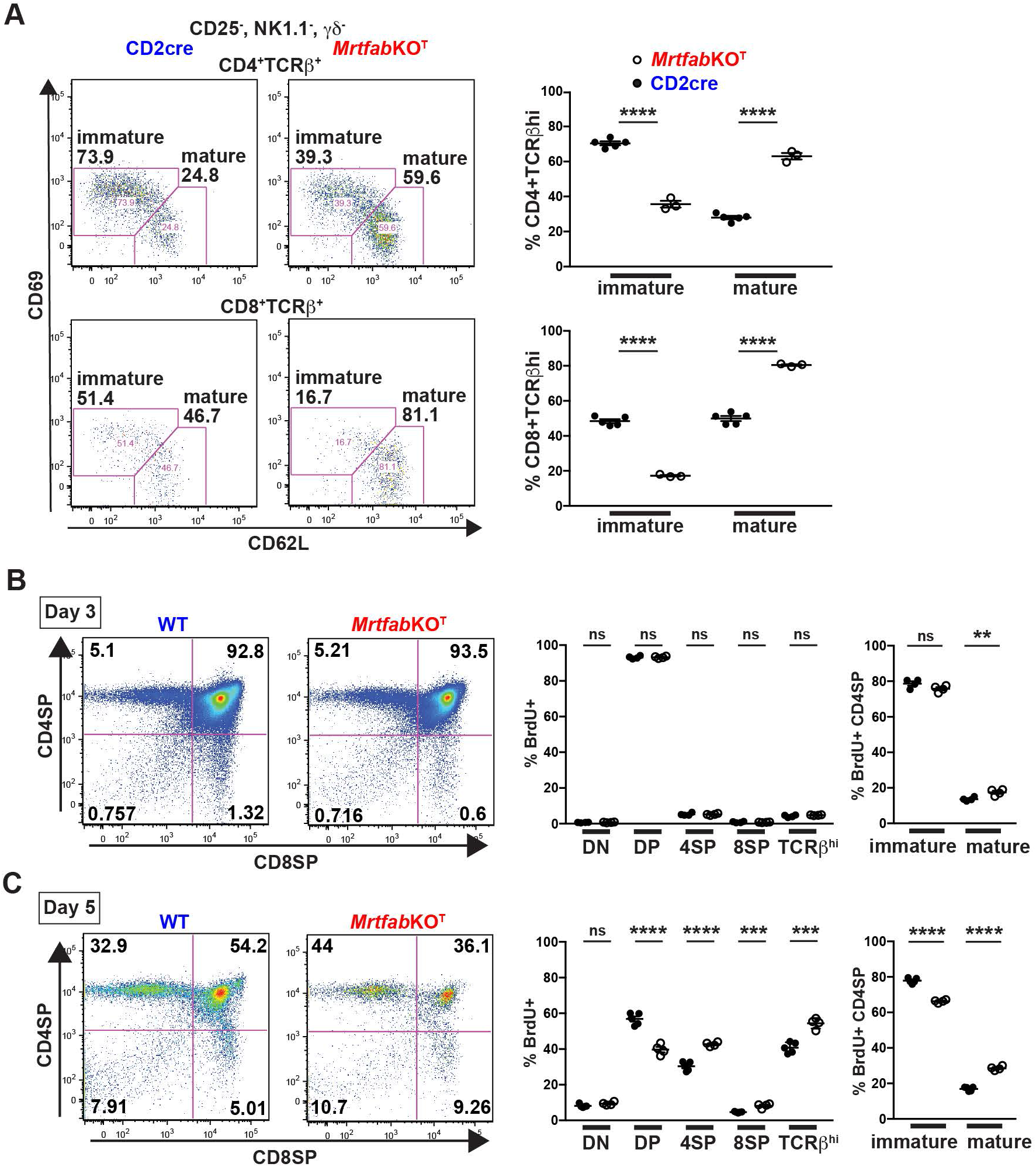
Mature thymocyte egress is impaired in *Mrtfab*KO^T^ mice. A. Representative CD69, CD62L profiles of live-gated, CD25-, GL3-NK1.1-CD4+ TCRβ and CD8+ TCRβ^hi^ thymocytes from age-matched female CD2-Cre and *Mrtfab*KO^T^ mice. Proportions of CD4+ TCRβ^hi^ and CD8+ TCRβ^hi^ cell are quantified at right. Data are combined from two independent experiments. B. WT and *Mrtfab*KO^T^ animals were injected with BrdU, and thymocytes analysed 3 days later. Left, representative BrdU+ gated CD4 and CD8 profiles. Centre, proportion of BrdU+ cells in DN, DP, CD4SP, CD8SP and TCRβhi populations. Right, proportions of BrdU+ thymocytes in immature CD69^hi^ CD62^lo^ and mature CD69^lo^ CD62L^hi^ TCRβ^hi^ CD4SP populations. 4 mice injected per genotype ^C.^ As in (B), with analysis 5 days after BrdU injection. 5 WT, 4 *Mrtfab*KO^T^ mice.

The increased numbers of mature SP cells seen in the MRTF-A/B null thymus could in principle arise through increased proliferation of the mature thymocyte population, increased supply of immature SP thymocytes from the DP pool, more rapid maturation of the newly selected SP cells, or defective thymic egress of the mature SPs. To examine these possibilities, we performed BrdU pulse-labelling experiments. Most thymocyte proliferation occurs during the DN and DP stages prior to TCRβ rearrangement and positive selection, with mature SP thymocytes exiting the thymus synchronously 4–5 d after selection (McCaughtry et al., 2007). In agreement with this, mature MRTF-null SP thymocytes from animals did not display appreciably more BrdU incorporation immediately after labelling, and three days post labelling, most DP cells were labelled (Figure 2B, Figure S2E). At this time, there were comparable small numbers of total labelled wildtype and MRTF-null SP thymocytes, indicating that their rates of exit from the DP pool were similar (Figure 2B). By 5 days after labelling, the overall proportion of BrdU-labelled thymocytes had substantially declined, and reflecting thymocyte differentiation and thymic exit; at this point the proportion of labelled MRTF-null SP thymocytes was increased relative to control, and the proportion of mature TCRβ^hi^ CD69^hi^ CD62L^lo^ SP cells increased (Figure 2C). Taken together these results suggest that the presence of increased numbers of mature SP cells in the MRTF-A/B null thymus reflects a reduced rate of thymic exit.

### MRTF inactivation disrupts thymic architecture and impairs cell migration

Previous studies have shown that defects in thymocyte egress and lymphocyte trafficking are associated with disruption of thymic architecture and impaired central tolerance (Mou, 2012 #8, for review see Cosway et al., 2021; Misslitz et al., 2004; Nitta et al., 2009). Hematoxylin and eosin (H&E) staining of MRTF-null thymus revealed dramatic disruption of the darker-staining cortex and the lighter-staining medulla, with extensive dark staining penetrating deep into the organ, and lighter staining regions greatly reduced and fragmented (Figure 3A). Staining for the endothelial marker CD31/PECAM1 and the stromal fibroblast marker ER-TR7 revealed that the perivascular spaces adjacent to the endothelial vessels were distended, containing multiple layers of CD3+ cells, in contrast to control CD2cre organs, in which a single layer of CD3+ cells was flattened against the vessel wall (Figure 3B).

**Figure 3.**
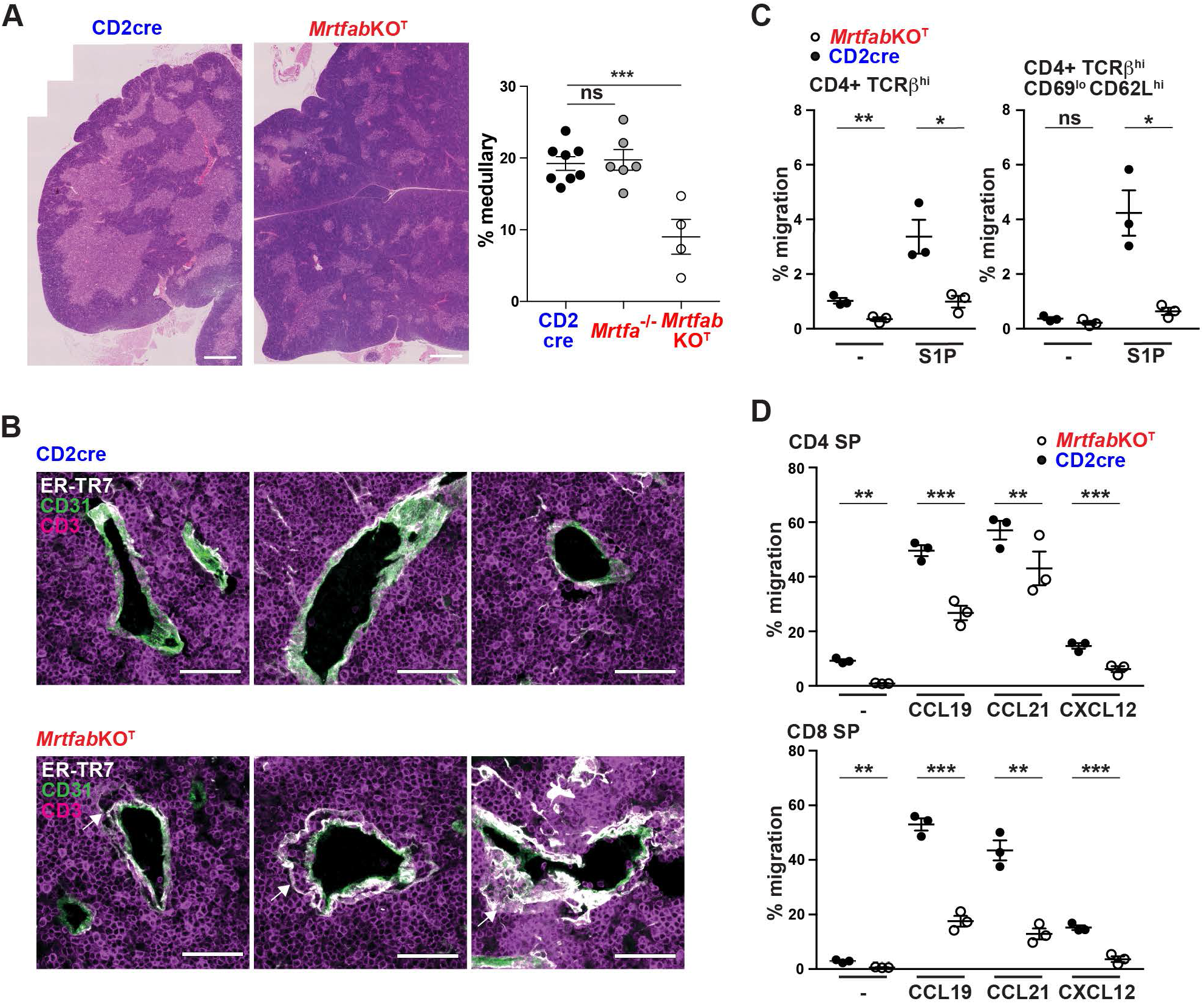
Aberrant morphology of MRTF-null thymus is associated with thymocyte migration defects. A. Representative hematoxylin and eosin stained sections of young adult female CD2cre and *Mrtfab*KO^T^ thymuses. Cortical regions stain dark, and medullary regions stain light. Scale bar, 500μm. Right, relative proportion of medullary areas, measured from 2-3 sections 250 μm apart, in *Mrtfab*KO^T^ and *Mrtfa*^-/-^ thymus. B. Immunofluorescence staining of the perivascular space in CD2cre and *Mrtfab*KO^T^ thymuses for the endothelial marker CD31/PECAM1. Scale bar, 50μm. Mature *Mrtfab*KO^T^ CD4+ thymocytes exhibit defects in S1P-dependent migration. C. Thymocytes from CD2cre and *Mrtfab*KO^T^ animals were allowed to migrate to 10nM S1P in transwells for 180 min, stained for CD4 and maturity markers, and compared to the input population. A representative of two independent experiments is shown; data are technical replicates. D. *Mrtfab*KO^T^ SP thymocytes exhibit defects in cytokine-dependent migration. Thymocytes from CD2cre and *Mrtfab*KO^T^ animals were allowed to migrate to CCL19, CCL21 or SDF-1 in transwells for 120 min, stained for CD4 and CD8, and compared to the input population. A representative of three independent experiments is shown; data are technical replicates.

MRTF-SRF signalling supports effective operation of cytoskeletal dynamics (for review see Olson and Nordheim, 2010) and we previously showed that is also true of hematopoietic and T cell lineages (Costello et al., 2015; Maurice et al., 2024). We therefore examined thymocyte responses to CCL19, CCL21, CXCL12, and S1P, the major mediators of thymocyte homing, intrathymic trafficking, and thymic egress. Expression of the CCL19/21 receptor CCR7 was slightly reduced in MRTF-null cells, while that of the CXCR4, the receptor for CXCL12 was unaffected (Figure S3A). Expression of S1PR1, the major receptor for S1P, which promotes thymocyte egress and is high in mature thymocytes, was further elevated in MRTF-null cells (Figure S3B). MRTF-null thymocytes exhibited defective migration towards CCL19, CCL21, CXCL12, and S1P in transwell assays (Figure 3C, 3D). Induction of F-actin assembly is central to the cellular response to migratory cues. MRTF-null thymocytes had lower basal F-actin levels than CD2cre controls, although S1P and chemokine stimulation remained able to induce F-actin assembly in both CD8+ and CD4+ thymocytes (Figure S3D). Moreover, levels of both β and γ actin mRNA were also significantly reduced in MRTF-null cells, as we previously observed in T-cells (Figure S3E). Taken together these data support the view that MRTF-SRF signalling supports cytoskeletal processes required for chemokine-mediated intra-thymic migration and S1P mediated thymic egress.

### *Mrtfab*KO^T^ animals are lymphopenic

Thymic egress, as well as proliferation of cells in the periphery, contributes to T cell homeostasis, so we next analysed peripheral T cells. Blood from *Mrtfab*KO^T^ animals was severely depleted in circulating TCRβ^hi^ lymphocytes, the CD4 and CD8 TCRβ^hi^ populations being reduced 2-4 fold (Figure 4A). T cell numbers in *Mrtfab*KO^T^ spleen and axillary, inguinal and mesenteric lymph nodes were also substantially reduced (Figure 4B); thus, the similar proportions of CD4 FoxP3+ T_reg_ cells present in wildtype and *Mrtfab*KO spleen and lymph nodes indicates that total T_reg_ numbers must be reduced in these organs (Figure 4C, Figure S4A, S4B). *Mrtfab*KO^T^ spleen and lymph node T cells also displayed elevated proportions of CD44^hi^ CD62L^lo^ cells in the absence of explicit antigenic stimulation (Figure 4D), a phenotype associated with lymphopenia-induced homeostatic proliferation (LIP, Goldrath et al., 2000).

**Figure 4.**
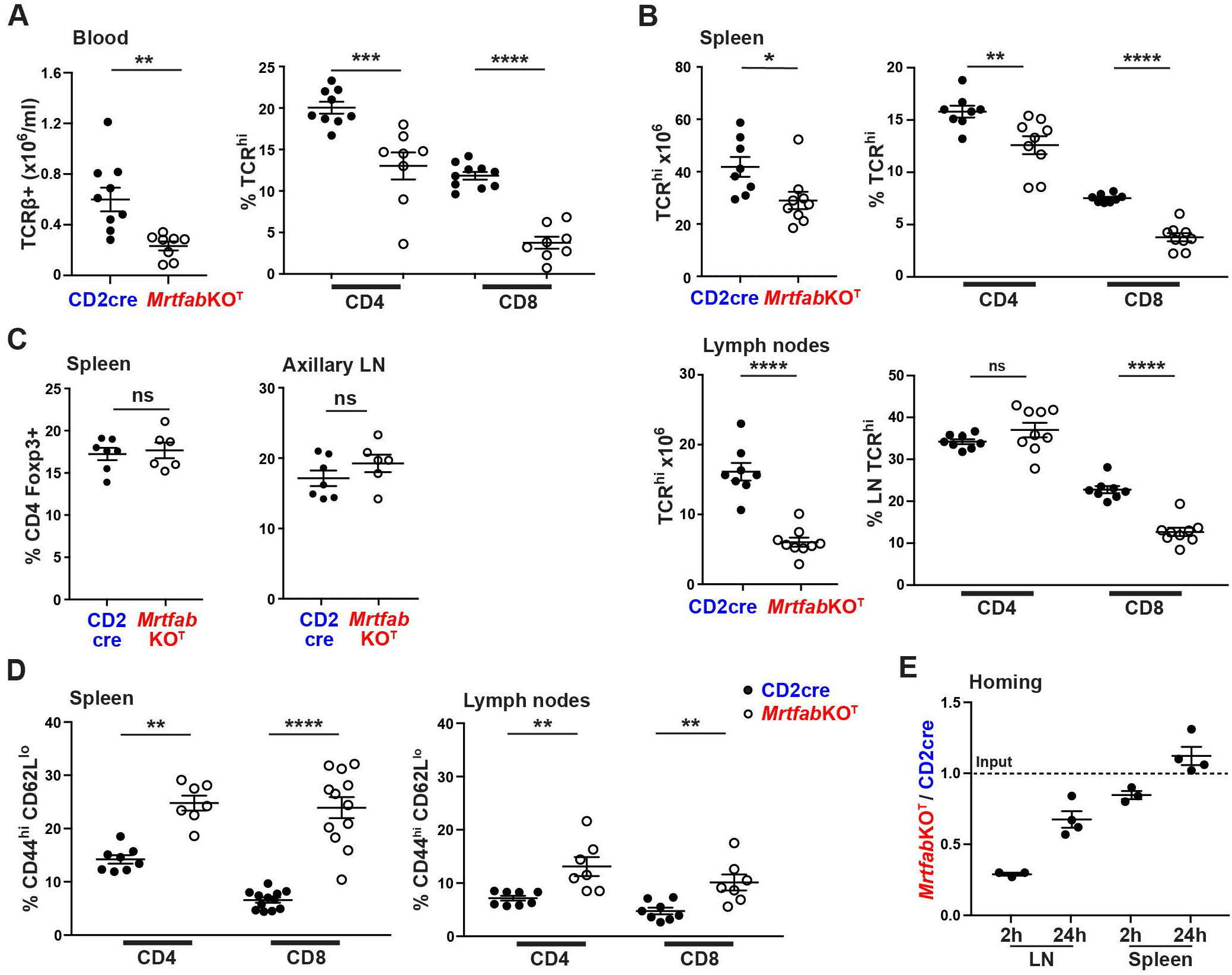
*MrtfabKO*^T^ mice are lymphopenic. A. Tail vein bleeds from CD2cre and *Mrtfab*KO^T^ mice were stained for CD4, CD8 and TCRβ. Data are total TCRβ+ cell numbers and proportions of CD4+ TCRβ+ and CD8 TCRβ+ cells, from two independent experiments, each with ≥2 female age-matched mice per genotype. B. Cells from spleen (upper panels) or pooled inguinal, axillary, brachial and mesenteric lymph nodes (lower panels) from CD2cre and *Mrtfab*KO^T^ mice were stained as in (A). Data are total TCRβ^hi^ cell numbers and proportions of CD4+ TCRβ^hi^ and CD8 TCRβ^hi^ cells from two independent experiments, each with ≥3 age-matched mice per genotype. C. Cells isolated from the spleen and axillary lymph nodes were surface stained with CD4, CD8, fixed and permeabilised for Foxp3 staining. Data show the percentage of Foxp3+ cells within the CD4+ gate. Data from 7 adult and 6 adult *Mrtfab*KO mice, combined from two independent experiments. D. Cells isolated from spleen and lymph nodes were stained with CD4, CD8, CD44 and CD62. Data show proportions of CD44^hi^CD62L^lo^ T cells within the CD4+ and CD8+ populations. Data derived from young adults collected over four independent experiments. E. CD2cre and *Mrtfab*KO^T^ T cells pooled from 5 animals of each genotype were labelled with CFSE and SNARF respectively and co-transferred into wildtype hosts at 1:1 ratio. Data are ratio ± SEM of *Mrtfab*KO^T^ to CD2cre cells in lymph nodes and spleen in each of 3 or 4 recipients at 2h and 24h following transfer, and are representative of two independent experiments.

We examined the possibility that these phenotypes arise from defective proliferation of peripheral T cells. Indeed, MRTF-null CD4 and CD8 T cells exhibited striking proliferative defects upon adoptive transfer into *Rag2*^-/-^ hosts (Figure S4C, S4D). This did not appear to result from intrinsic defects in the ability to proliferate *per se*, however, because MRTF-null T cells proliferated normally in response to TCR crosslinking (Figure S4E, see also Maurice et al., 2024), remained responsive to the homeostatic cytokine IL-15 *in vitro* (Figure S4F), and maintained their survival response to IL-7 (Figure S4G).

Previous studies have shown that homeostatic proliferation is critically dependent on recirculation of lymphocytes via lymph nodes (Schuster et al., 2009). Moreover, the preceding results show that MRTF-null thymocytes are defective in migration *in vitro*, and in thymic homing and egress *in vivo*. We therefore assessed the homing of *Mrtfab*KO^T^ T cells to peripheral lymphoid organs by injecting labelled wildtype and *Mrtfab*KO^T^ CD8 T cells into the tail vein of wildtype mice. *Mrtfab*KO^T^ T cell numbers in the inguinal lymph node were greatly reduced at 2h after injection, recovering after 24h; in contrast, MRTF-A/B null T cell homing to the spleen was barely affected (Figure 4E). Defects in T cell homing to lymph nodes but not spleen also results from defective CCR7 signalling (Forster et al., 1999). These results suggest that inefficient homing to lymph nodes contributes to the lymphopenic phenotype of *Mrtfab*KO^T^ animals. Reduced total T_reg_ numbers are likely to further exacerbate LIP by diminishing regulation of APC activation and T cell expansion (Picca et al., 2006; Winstead et al., 2008).

### *Mrtfab*KO^T^ mice exhibit signs of defective tolerance

Previous studies have demonstrated that disrupted thymic cortical medullary structure and defects in CCR7-mediated intra-thymic trafficking are associated with breakdown in central tolerance (Nitta, 2009 #78, for reviews see Cosway et al., 2021; Forster et al., 2008; James et al., 2018). To address this, we examined tissues in *Mrtfab*KO^T^ mice for evidence of lymphocyte infiltration and inflammation. The lacrimal glands of young *Mrtfab*KO^T^ mice exhibited extensive lymphocyte infiltration, including CD4+, CD8+ and FoxP3+ T cells, and inflammation (Figure 5A, 5B, Figure S5A). This appeared more pronounced in older animals, which exhibited severe tissue damage, with evidence of tertiary lymphoid structures with extensive B cell infiltration (Figure S5B-D). Both male and female *Mrtfab*KO^T^ mice also weighed significantly less than age-matched CD2cre controls (Figure 5C). Age-matched cohorts of ∼13 and ∼21 week old *Mrtfab*KO^T^ mice exhibited elevated levels of fecal lipocalin 2 (LCN2), a non-invasive biomarker of intestinal inflammation (Figure 5D). Sectioning of the large intestine revealed areas of substantial hyperplasia, accompanied by T cell infiltrates, a phenotype which was particularly pronounced in males (Figure 5E, 5F). Transfer of *Mrtfab*KO^T^, but not wildtype CD2cre thymocytes into *Rag2*^-/-^ recipients, which lack T and B cells, also induced lacrimal exocrinopathy, indicating that it reflects an intrinsic defect in MRTF-null thymocytes (Figure S5E, S5F). Taken together these results indicate that *Mrtfab*KO^T^ animals are defective in central tolerance.

**Figure 5.**
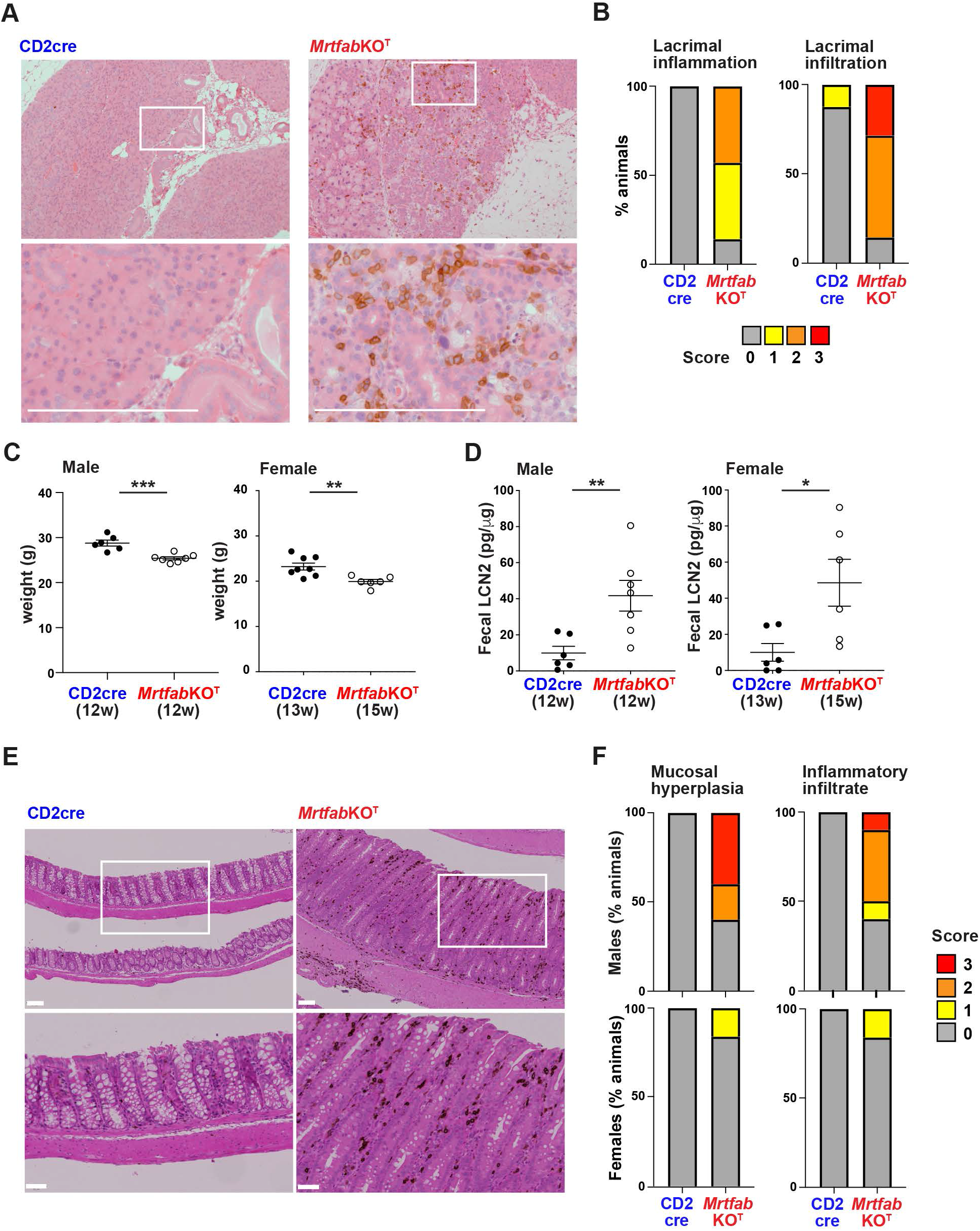
*MrtfabKO*^T^ mice exhibit autoimmune phenotypes. A. Images of 3μm lacrimal gland sections from CD2Cre and *Mrtfab*KO^T^ mice (median ages 9 and 10 weeks respectively), stained with hematoxylin and eosin, and anti-CD3 (brown). Boxed sections are enlarged below. Scale bar, 1mm. The *Mrtfab*KO^T^ image is from animal graded as moderate for inflammation and infiltration in (B). B. Qualitative grading of lacrimal gland inflammation and infiltration in young CD2Cre (n=8, 5 female, 3 male) and *Mrtfab*KO^T^ (n=7, 4 female, 3 male) mice as in (A). C. Weights of age- and sex-matched CD2Cre and *Mrtfab*KO^T^ mice. Median ages are given in weeks. Data points are mean ± SEM. D. Fecal lipocalin 2 levels in the mice analysed in (C). Data points are mean ± SEM. E. Mucosal hyperplasia in the large intestine of 30 week old male *Mrtfab*KO^T^ mice. Upper panels, 3μm sections of large intestine stained with hematoxylin and eosin, and anti-CD3 (brown); scale bars, 100μm. Lower panels, enlargements of the boxed areas; scale bars, 50μm. F. Qualitative grading of mucosal inflammation (left) and inflammatory infiltration (right) in age-matched males (top; CD2cre, n = 16, median age 33w; *Mrtfab*KO^T^, n = 10; median age, 30w) and females (bottom CD2cre, n = 8, median age, 31w; *Mrtfab*KO^T^, n = 6; median age, 31w).

### Negative selection is intact in MRTF-null thymocytes

We next investigated the basis for the autoimmune defect seen in *Mrtfab*KO^T^ animals. First We first analysed negative selection, because in CCR7R-null animals related phenotypes are associated with defective negative selection (Nitta et al., 2009), and thymic egress timing underlies negative selection of self-reactive CD8+ thymocytes (Badr et al., 2023). Irradiated wildtype or RIP-mOva CD45.1/CD45.2hosts were reconstituted with bone marrow carrying the mOva-specific OT-I or OT-II TCR transgenes (Barnden et al., 1998; Hogquist et al., 1994). Analysis of thymocytes 6 weeks post-reconstitution with OT-I TCR+ bone marrow revealed efficient selection of CD2cre and *Mrtfab*KO^T^ CD8+ SP thymocytes in wildtype hosts (Figure S6A) and positively selected OT-I thymocytes were enriched in mature vα2+ CD24^lo^ cells, as seen with the polyclonal populations (Figure 6A). In contrast, both wildtype and *Mrtfab*KO^T^ OT-I thymocytes were efficiently deleted upon reconstitution of RIP-mOva+ hosts (Figure 6B, Figure S6B). Virtually complete deletion of both wildtype and *Mrtfab*KO^T^ cells was apparent upon analysis of blood, spleen and lymph nodes (Figure 6C). Similar results were obtained the same approach was used to analyse negative selection of cells expressing the MHCII-restricted OT-II TCR (Figure 6D-F; Figure S6B). OT-II thymocyte deletion also occurred efficiently in primary CD2cre OT-II or *Mrtfab*KO^T^ RIP-mOva animals (Figure 6D-F; Figure S6C-E). Thus, negative selection is intact in *Mrtfab*KO^T^ animals.

**Figure 6.**
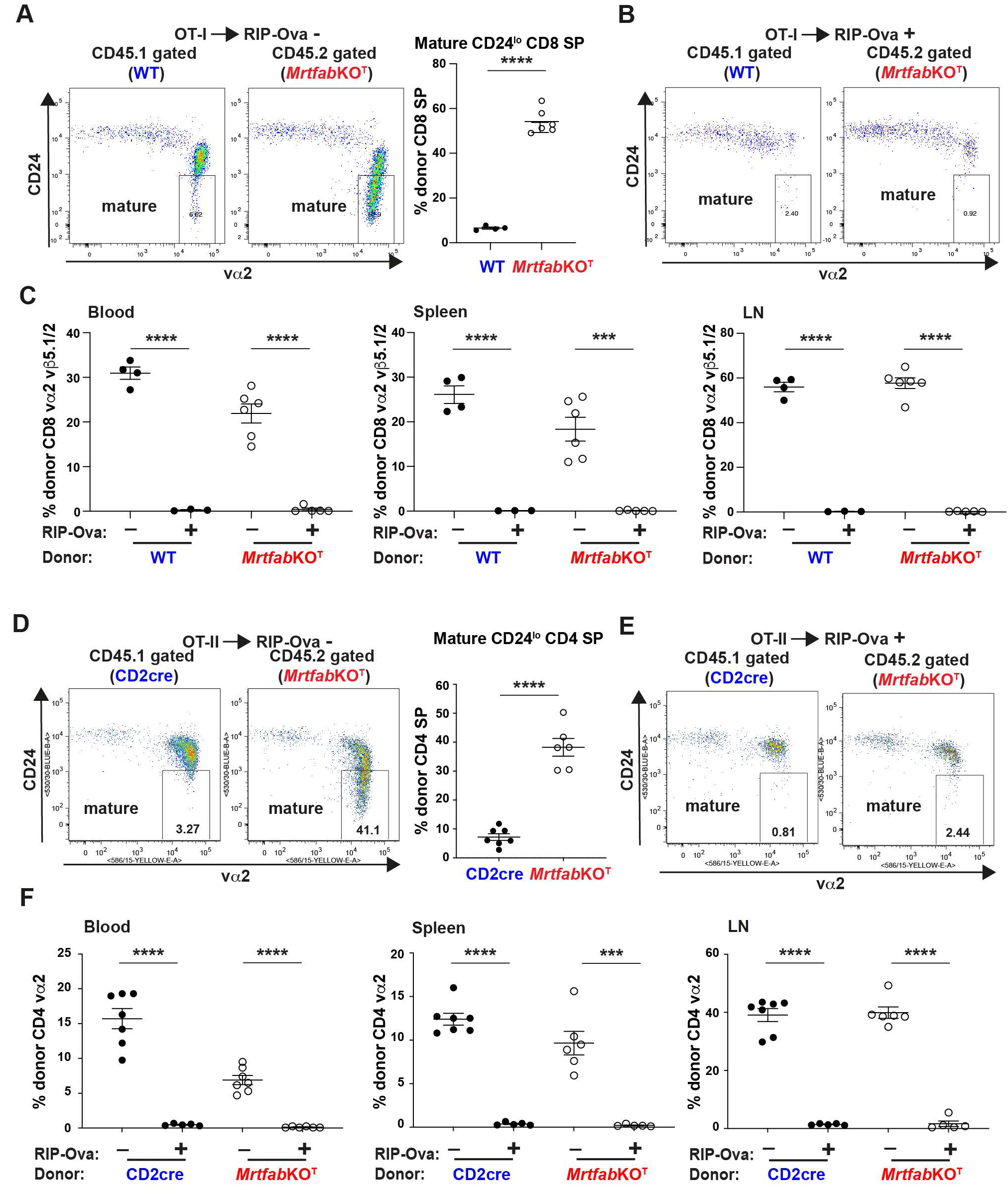
Negative selection is intact in *MrtfabKO*^T^ mice. A. Separate cohorts of irradiated 10 week old female wildtype CD45.1/CD45.2 hosts were reconstituted with bone marrow from either wildtype (CD45.1) or *Mrtfab*KO^T^ (CD45.2) OT-I TCR transgenic animals. Left, representative CD24 / vα2 profiles of CD8+ thymocytes from OT-I WT and OT-I *Mrtfab*KO^T^ animals, gated on CD45.1 and CD45.2 respectively. Right, proportions of mature CD24^lo^ thymocytes in the two populations. Data are mean ± SEM, combined from two independent experiments (WT, n=4; *Mrtfab*KO^T^, n=6). For CD4/CD8 profiles see Figure S6A. Primary animal data are shown in Figure S5C. B. Separate cohorts of irradiated female RIP-mOva+ CD45.1/CD45.2 hosts were reconstituted with wildtype (CD45.1) or *Mrtfab*KO^T^ (CD45.2) OT-I TCR transgenic bone marrow, and analysed as in (A). Data are representative of two independent experiments (WT, n=3; *Mrtfab*KO^T^, n=5). For CD4/CD8 profiles see Figure S6B. C. Proportions of donor-gated CD8+ vα2+ vβ5.1/2+ T cells in peripheral blood, spleen and lymph nodes. Data are mean ± SEM, combined from the experiments illustrated in (A) and (B). D. Wildtype CD45.1/CD45.2 hosts were reconstituted with bone marrow from either CD2cre (CD45.1) or *Mrtfab*KO^T^ (CD45.1) OT-II TCR transgenic animals as in (A). Left, representative CD24 / vα2 profiles of CD4+ thymocytes. Right, proportions of mature CD24^lo^ thymocytes in the two populations. Data are mean ± SEM, combined from two independent experiments (CD2cre, n=7; *Mrtfab*KO^T^, n=6). E. Irradiated female RIP-mOva+ CD45.1/CD45.2 hosts were reconstituted with CD2cre (CD45.1) or *Mrtfab*KO^T^ (CD45.2) OT-II TCR transgenic bone marrow, and analysed as in (D). Data are representative of two independent experiments (CD2cre, n=5; *Mrtfab*KO^T^, n=6). F. Proportions of donor gated CD4+ vα2+ T cells in peripheral blood, spleen and lymph nodes. Data are mean ± SEM, combined from the experiments illustrated in (D) and (E).

### MRTF-null T_reg_ cells home to sites of inflammation but are functionally defective

We next evaluated the role of T_reg_ cells in the autoimmune pathology of *Mrtfab*KO^T^ animals. Pilot experiments indicated the lacrimal gland pathology induced in *Rag2*^-/-^ animals by *Mrtfab*KO^T^ thymocyte transfer could be suppressed by co-transfer of wildtype CD2cre T_reg_ cells (Figure S6F), suggesting that *Mrtfab*KO^T^ T_reg_ cells are defective. To investigate this in more detail, we exploited the observation that transfer of conventional CD4+ T cells into immunodeficient hosts induces intestinal hyperplasia and lymphocyte infiltration that can be suppressed by co-transferred T_reg_ cells (Powrie et al., 1993).

Transfer of 300,000 purified wildtype CD4+CD25-CD45RB^hi^ T_con_ cells into *Rag2*^-/-^ recipients led to decreased weight gain, and/or spontaneous sustained weight loss that necessitated culling under our animal procedure guidelines (Figure 7A, Figure S7A). At time of cull, animals displayed severe intestinal pathology with hyperplasia and lymphocyte infiltration (Figure 7B). Co-transfer of 150,000 CD4+CD25+CD45RB^lo^ T_reg_ cells purified from CD2cre animals suppressed both the mortality and the intestinal pathology induced by T_con_ cell transfer, but co-transfer of *Mrtfab*KO^T^ T_reg_ cells did not (Figure 7D), even though FoxP3+ T_reg_ cells were detectable in the intestine (Figure 7C,7D; Figure S7B). The intestinal pathology was accompanied by increased cellularity in the mesenteric lymph nodes (MLNs), but not the spleen, which was effectively suppressed by co-transfer of wildtype T_reg_ cells (Figure 7E). In contrast, the increased MLN cellularity was unaffected by cotransfer of *Mrtfab*KO^T^ T_reg_ cells, although FoxP3+ cells were readily detectable in MLN, indicating that their homing to peripheral lymphoid organs was intact (Figure 7E). Thus, *Mrtfab*KO^T^ T_reg_ are functionally defective *in vivo*.

**Figure 7.**
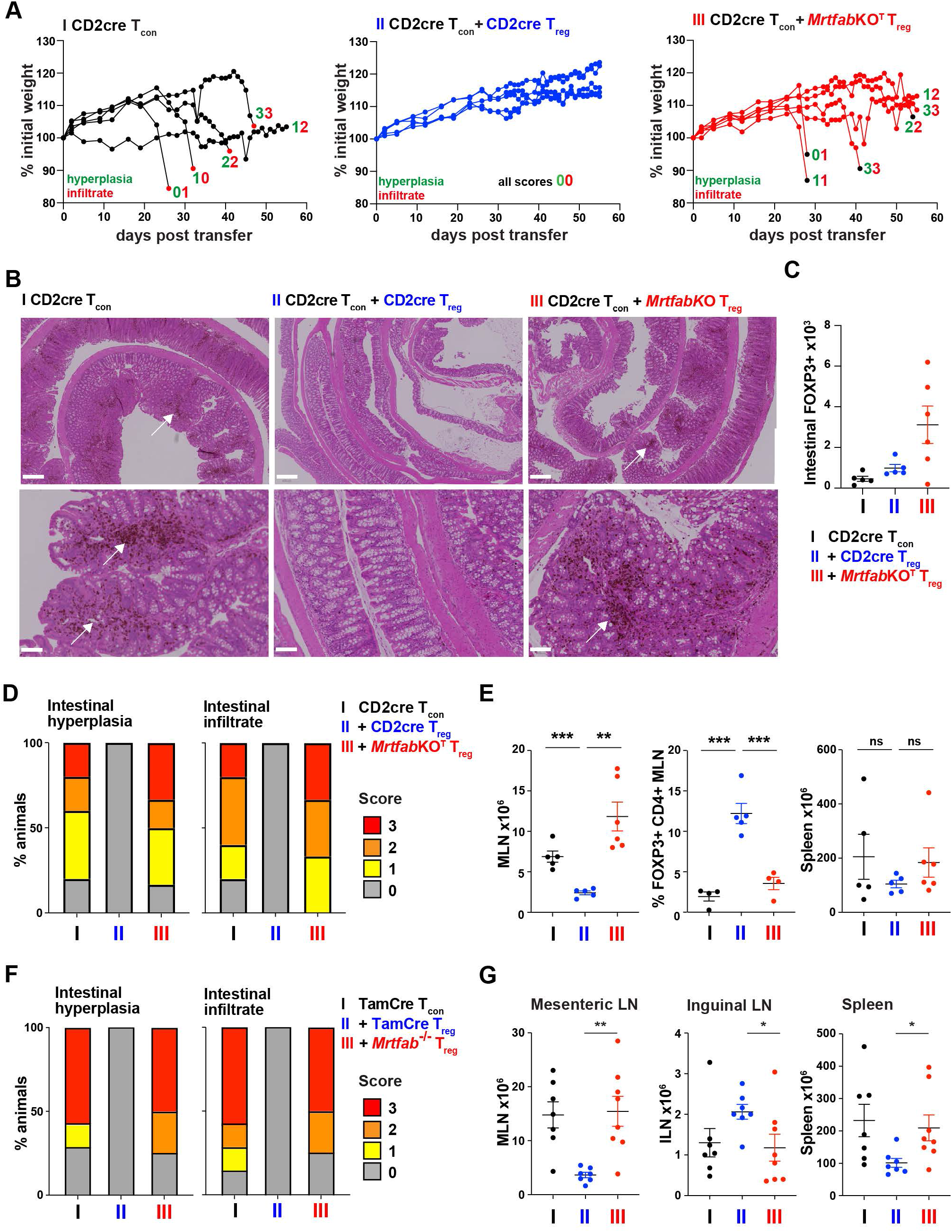
*MrtfabKO*^T^ T_reg_ cells are functionally deficient *in vivo* A. *In vivo* T_reg_ suppression assay. Mortality following transfer of 300,000 CD2cre T_con_ (CD4+ CD25-CD45RB^hi^) cells, either alone or with 150,000 CD2cre or *Mrtfab*KO^T^ T_reg_ (CD4+ CD25+ CD45RB^lo^) cells into male *Rag2*^-/-^recipients. Animals were culled upon loss of >15% starting weight or persistent loss of condition. Hyperplasia or infiltrate pathology scores at cull point or end of experiment are indicated. A similar but less pronounced result was obtained using female hosts. B. Large intestinal hyperplasia and lymphocyte infiltration (brown, arrows) following T_con_/T_reg_ transfer as in (A). Left, 10x images of representative 3μm large intestine sections stained with hematoxylin and eosin, and anti-CD3 (brown), made at point of cull or 10 weeks after transfer. Scale bars, 400μm (upper panels), 100μm (lower panels). C. Numbers of FoxP3+ cells in large intestine made at point of cull or 10 days following T_con_/T_reg_ transfer as in (A). For images, see Figure S7B. D. Qualitative grading of large intestine hyperplasia and infiltration in the experiment shown in (A). WT T_con_ alone, n=5; WT Tc_on_ + CD2cre T_reg_, n=5; WT Tc_on_ + *Mrtfab*KO^T^ T_reg_, n=6. E. Left, mesenteric lymph node cellularity and proportion of FoxP3+ CD4+ T cells; right splenic cellularity in the experiment shown in (A). Data are mean ± SEM, WT T_con_ alone, n=5; WT Tc_on_ + CD2cre T_reg_, n=5; WT Tc_on_ + *Mrtfab*KO^T^ T_reg_, n=6. F. Defective suppression by *Mrtfab*KO^T^ T_reg_ cells does not reflect a developmental defect. Male *Rag1^-^*^/-^ recipients were injected with 300,000 TamCre T_con_ (CD4+ CD25-CD45RB^hi^) cells, either alone or with 150,000 TamCre or *Mrtfab*^-/-^, T_reg_ (CD4+ CD25+ CD45RB^lo^) cells, generated by peripheral inactivation of *Mrtfb* in *Mrtfa*^-/-^ *Mrtfb*^fl/fl^ cells using Cre^ERT^ (Maurice et al., 2024). Qualitative grading of large intestine hyperplasia and infiltration is summarised. For representative images see Figure S7F. WT T_con_, n=10; WT Tc_on_ + CD2cre, n=8; WT Tc_on_ + *Mrtfab*KO^T^, n=7. A representative of two independent experiments, with similar results, is shown. G. Cellularity of peripheral lymphoid organs in the suppression experiment shown in (F).

To address whether the *Mrtfab*KO^T^ T_reg_ defect results from aberrant T_reg_ development, we induced MRTF inactivation in mature peripheral T-cells (Maurice et al., 2024). Rag2^-/-^ mice were reconstituted with bone marrow from TamCre *Mrtfa*^-/-^*Mrtfb*^fl/fl^ or TamCre animals, fed 4-hydroxytamoxifen (4OHT) to inactivate Mrtfb in susceptible cells, and the resulting mature TamCre *Mrtfab*^-/-^ or TamCre control CD4+CD25+CD45RB^lo^ T_reg_ cells purified for use in transfer experiments (Figure S7C). Transfer of 300,000 purified TamCre T_con_ cells into *Rag1*^-/-^ recipients again led to decreased weight gain, and/or spontaneous sustained weight loss, associated with intestinal and lacrimal gland pathology (Figure 7F; Figure S7D-F). Both could be suppressed by co-transfer of 150,000 purified TamCre T_reg_ cells, but not by TamCre *Mrtfab*^-/-^ T_reg_ cells (Figure 7F; Figure S7D-F). Intestinal pathology was again accompanied by increased mesenteric lymph node cellularity, and this was also suppressed by TamCre T_reg_ cells but not by TamCre *Mrtfab*^-/-^ T_reg_ cells (Figure 7G, Figure S7G). These data show that *Mrtfab*KO^T^ CD4+CD25+CD45RB^lo^ T_reg_ cells develop normally but are functionally defective *in vivo*. *Mrtfab*KO^T^ T_reg_ cells nevertheless behaved similarly to wildtype cells *in vitro*, however. Upon TCR crosslinking, they actually overproduced the suppressor cytokine IL-10 (Figure S7H), while upon presentation of Ova peptide by DC, they could suppress both CD44 and CD25 upregulation and proliferation of co-cultured CD4+ OT-II T cells (Figure S7I).

Taken together with the results in the preceding sections, we data show that MRTF-SRF signalling is required for central tolerance. This reflects a role in T_reg_ function rather than T_reg_ development, and appears to affect the suppressive function of T_reg_ cells *in vivo*, given that *Mrtfab*KO^T^ cells remain competent to mount a suppressive response *in vitro*.

## DISCUSSION

Here we have demonstrated that the actin-regulated MRTFs, transcriptional coactivators of the SRF transcription factor, play important roles in cell migratory events during thymocyte development, T cell trafficking and homeostasis. MRTF activity is required for effective homing of early thymocyte precursors, efficient thymic egress, T cell homing to secondary lymphoid organs, and maintenance of thymic architecture. Animals with MRTF-null T cells are also lymphopenic, and exhibit defects central tolerance: while negative selection is intact, MRTF-null T_reg_ cells are unable to suppress inflammation and lymphocyte infiltration in lacrimal gland and intestinal inflammation models. MRTF-null phenotypes are similar in many ways to those arising from inactivation of CCR7- and S1P-dependent signal pathways, and MRTF null thymocytes and T cells exhibit defects in F-actin assembly and migratory responses to multiple migratory stimuli, including sphingosine 1-phosphate (S1P), CCR7 ligands, and SDF-1. MRTF inactivation phenotypes are also similar to those seen upon inactivation of numerous mediators of cytoskeletal dynamics. Our results thus show that MRTF-SRF signalling is required for the execution of signal-induced cytoskeletal changes rather than function of specific receptors, consistent with its role in other settings.

Our MRTF inactivation model relies on the conditional inactivation of *Mrftb* in Mrtfa^-/-^ animals (Mokalled et al., 2015) using hCD2icre, thereby generating thymocytes that are MRTF-null (*Mrtfab*KO^T^) from the DN2/3 stage onwards. We did not address functional redundancy between the MRTFs, but previous studies by us and others indicate that they function redundantly in both T cells and other cell types (Costello et al., 2015; Mokalled et al., 2015). The MRTFs are not required for the early stages of thymocyte development and proliferation, and thymocyte positive and negative selection both proceeded normally, as did generation of CD4 FoxP3+ T_reg_ cells. However, MRTF inactivation did result in increased numbers of both thymic γδ T cells and mature *Mrtfab*KO^T^ thymocytes; this is also likely to reflect MRTF-dependent migration defects, as discussed below. We previously showed that although TCF-SRF signalling is necessary and sufficient for positive selection, animals lacking all three TCFs nevertheless produce reduced numbers of SP thymocytes and T cells (Costello et al., 2010; Costello et al., 2004; Mylona et al., 2011). It therefore remains possible that the MRTFs can functionally substitute for the TCFs in positive selection, even though MRTF-SRF signalling plays no essential role.

Our results show that the MRTFs are required for effective thymic egress, and that MRTF inactivation resulted in an increased accumulation of mature SP thymocytes within a distended perivascular space. Egress of mature SP thymocytes is dependent on signalling by sphingosine 1-phosphate receptor S1PR1 (Allende et al., 2004; Matloubian, 2004 #37, for review see Cyster and Schwab, 2012), on KLF2, the factor that controls S1PR1 expression (Bai et al., 2007; Carlson et al., 2006), and other genes controlling S1P synthesis and turnover (for review seeCyster and Schwab, 2012). Consistent with this, we found that *in vitro Mrtfab*KO^T^ thymocytes exhibit defective chemotactic and cytoskeletal responses to S1P stimulation even though S1PR1 expression is elevated. It is likely that defective thymic egress accounts for the elevated γδ cell numbers seen in *Mrtfab*KO^T^ animals a phenotype also associated with inactivation of S1PR1 and KLF2 (Odumade et al., 2010). Thymic egress defects can also result from inactivation of general mediators of thymocyte and T cell adhesion and migration, including the LFA-1 integrin (Bose et al., 2014), Mst1/2 (Dong et al., 2009; Mou et al., 2012) and their effector kinases Ndr1/2 (Tang et al., 2015), and the actin regulators coronin 1a (Shiow et al., 2008) and mDia (Sakata et al., 2007). These results thus provide additional support for the view that MRTF-SRF signalling is a prerequisite for effective cytoskeletal regulation in diverse settings, including T cells (reviewed by Maurice et al., 2024; Olson and Nordheim, 2010, see).

MRTF inactivation in thymocytes caused disruption of thymic architecture, with the proportion of cortex increasing and medulla areas becoming fragmented. Reduced and fragmented thymic medulla is also seen upon inactivation of CCR7 or its ligands (Misslitz et al., 2004; Ueno et al., 2002). This is thought to reflect disruption of cortex-medulla migration, which is strongly CCR7-dependent (Kurobe et al., 2006), and is consistent with the defective cytoskeletal responses of *Mrtfab*KO^T^ thymocytes and T cells to the CCR7 ligands CCL19 and CCL21. In contrast, inactivation of the S1P receptor increases rather than decreases medullary area, presumably due to increased retention of mature thymocytes (Allende et al., 2004). Since MRTF inactivation impairs the responses to both S1P and CCR7 ligands, our data suggest the effect of MRTF inactivation on CCR7-dependent thymocyte medulla entry may outweigh any impairment of S1PR1-dependent thymic exit. Inactivation of the Mst1/2 kinases also reduces and fragments thymic medullary area (Mou et al., 2012), but effects of inactivating other cytoskeletal regulators have not been reported.

*Mrtfab*KO^T^ animals exhibit profound lymphopenia, with greatly reduced numbers of αβ T cells in the blood and in secondary lymphoid organs, including lymph nodes and spleen. This does not reflect cell-intrinsic defects, since *in vitro* Mrtf-null thymocytes and T cells exhibit normal proliferative and survival responses to TCR, IL-15 and IL-7 stimulation, while their initial proliferative response to TCR activation upon infectious challenge *in vivo* is also unimpaired (Maurice et al., 2024). Instead, we found that MRTF inactivation leads to defective homing to lymph nodes. Lymphopenia is also associated with defects in CCR7 and S1P receptors and their regulators (Forster et al., 1999)Allende, 2004 #38;Matloubian, 2004 #37;Bai, 2007 #63;Carlson, 2006 #62;Kuo, 1997 #112, for review see \Cyster, 2012 #61}, we think it like that. The lymphopenia seen in *Mrtfab*KO^T^ animals is thus likely at least in part to reflect defective cytoskeletal responses to CCR7 and S1P receptor-mediated signalling. Lymphopenia is associated with inactivation of several genes involved in T cell cytoskeletal dynamics, including LFA1, mDia1, Coronin, Mst1/2 and Ndr1/2 (Bose et al., 2014; Dong et al., 2009; Mou et al., 2012; Sakata et al., 2007; Shiow et al., 2008; Tang et al., 2015), and the MRTFs appear to fall into this category.

*Mrtfab*KO^T^ animals display signs of mucosal autommunity, with lacrimal and salivary gland inflammation and T-lymphocyte infiltration; they also weighed less than control animals and exhibited intestinal inflammation and infiltration. The exposure of mucosal surfaces are exposed to external antigens, commensal microbes, and pathogens might render them particularly be susceptible to immune pathology. The autoimmune phenotypes are reminiscent of but appear less severe that those seen in animals with defects in receptor-mediated thymocyte and T cell trafficking, including CCR7 and its ligands (Davalos-Misslitz et al., 2007; Hopken et al., 2007; Kocks et al., 2007; Kurobe et al., 2006; Nitta et al., 2009), KLF2, which plays important roles in controlling S1P signalling (Pabbisetty et al., 2016), and other regulators of thymocyte and T-cell motility such as Mst1 (Ueda et al., 2012).

Our data supports a model in which the autoimmunity seen in *Mrtfab*KO^T^ animals arises from defective T_reg_ function. The lacrimal gland phenotype was recapitulated upon *Mrtfab*KO^T^ thymocyte transfer into Rag2-/-hosts, and could be suppressed by co-transfer of wildtype T_reg_ cells, and in the Powrie experimental colitis model, intestinal mucosal inflammation could be suppressed by wildtype, but not by *Mrtfab*KO^T^ T_reg_ cells. In contrast, we found that negative selection of MHC class I- or class II-restricted transgenic TCRs against tissue-restricted antigens proceeds normally in *Mrtfab*KO^T^ animals. The migration defect in *Mrtfab*KO^T^ thymocytes thus not limiting for the early thymic exit recently reported to be required for elimination of self-reactive MHC class I-restricted CD8 T cells (Badr et al., 2023).

Functional defects in T_reg_ cells also underlie at least some of the CCR7/CCR7L null autoimmune phenotypes: wildtype but not CCR7/CCR7L null T_reg_ cells can suppress mucosal autoimmunity (Ishimaru et al., 2010) colitis in the Powrie colitis model (Schneider et al., 2007). We note, however, that the autoimmunity associated with CCR7/CCR7L deficiency is more severe than in *Mrtfab*KO^T^ mice (Davalos-Misslitz et al., 2007; Kurobe et al., 2006), and that CCR7/CCR7L-null animals also exhibit defective negative selection of TCRs against tissue-restricted antigens (Nitta et al., 2009; Ueno et al., 2004).

The nature of the defect in *Mrtfab*KO^T^ T_reg_ cells remains unclear. It cannot reflect a developmental defect in the TCR repertoire, because T_reg_ cells generated by peripheral inactivation of the MRTFs remained able to suppress colitis. *Mrtfab*KO^T^ T_reg_ cells exhibited normal cell surface marker profiles, and could suppress upregulation CD25 and CD44 on CD4 T cells normally upon cognate antigen presentation by DC *in vitro*. Moreover, although IL-10 plays a critical role in the suppressive effect of T_reg_ cells in the Powrie colitis model (Branchett et al., 2024), *Mrtfab*KO^T^ T_reg_ cells actually produced more IL-10 upon TCR crosslinking *in vitro*. Given the effects of MRTF inactivation on cytoskeletal dynamics and T-cell trafficking, we favour the view that *Mrtfab*KO^T^ T_reg_ cells have no intrinsic defects in their ability to respond to activation, but are instead unable to undertake cell trafficking or interactions required for effective suppression *in vivo*. *Mrtfab*KO^T^ T_reg_ cells were readily detectable in the inflamed intestine, but we note that intestinal homing is not required for their suppressive effect in the Powrie colitis model (Denning et al., 2005). CCR7-null T_reg_ cells have been reported to exhibit defects in lymph node homing and activation (Ishimaru et al., 2010; Kocks et al., 2007; Menning et al., 2007; Schneider et al., 2007), but we have seen no such phenotypes in polyclonal *Mrtrab*KO^T^ T_reg_ populations. We are currently using genetically labelled T_reg_ cells expression transgenic TCRs as a sensitive approach to tracking *Mrtrab*KO^T^ T_reg_ cell dynamics *in vivo*.

## ACKNOWLEDGEMENTS

We thank Jessica Diring for help with T-cell migration and imaging, and thank Carola Vinuesa, Gitta Stockinger and lab members for helpful discussions during the project and/or comments on the manuscript. We thank Bill Heath (WEHI, Melbourne, Australia) for OT-I TCR, OT-II TCR and *RIP-mOva* transgenic mice (Barnden et al., 1998; Kurts et al., 1996), and Alejandro Suárez-Bonnet and Eleanor Herbert (Crick Experimental Histopathology STP) for scoring the pathology slides, and Clare Watkins, Julie Bee and the Biological Resources Facility for expert support.

This work was supported by the Francis Crick Institute which receives its core funding from Cancer Research UK (CC2102), the UK Medical Research Council (CC2102), and the Wellcome Trust (CC2102). This research was funded in whole, or in part, by the Wellcome Trust CC2102. For the purpose of Open Access, the authors have applied a CC BY public copyright licence to any Author Accepted Manuscript version arising from this submission.

## AUTHOR CONTRIBUTIONS

PC conceived the project. PC, DM, and MB-J constructed mouse strains, and designed, performed and interpreted experiments; RT designed and interpreted experiments. PC and RT wrote the paper with PC and input from the other authors.

## METHODS

### Mice

For gene inactivation in the thymus we used CD2-iCre (de Boer et al., 2003), with *Mrtfa*^-/-^ and *Mrtfb*^fl/fl^ (Mokalled et al., 2010). Peripheral inactivation of *Mrtfb* was achieved by tamoxifen-mediated *Mrtfb*^fl/fl^ inactivation in *Mrtfa*^-/-^*Mrtfb*^fl/fl^R26CreERT2 (*tm9(cre/ESR1)Arte*) mice as described (Maurice et al., 2024). OT-I TCR, OT-II TCR and *RIP-mOva* transgenic mice were as described (Barnden et al., 1998; Kurts et al., 1996). *Rag1*^-/-^ or *Rag2^-/-^* mice were used for thymocyte transfer and *in-vivo* suppression assays as indicated in the text. Reconstitution into irradiated CD45.1/CD45.2 (Shen et al., 1985) or *RIP1-mOva* (Kurts et al., 1996) mice used for negative selection studies. Both male and female mice were used. Genotyping (Transnetyx) used KAPA 2G fast genotyping mix (KAPA Biosystems). Animals were maintained under specific-pathogen-free conditions in The Francis Crick Biological Resources Unit. Animal experimentation, approved by the Crick Animal Ethics committee, was carried out under Home Office licence PP0389970 and PP0792028. Where relevant, age and/or sex of mice used is specified in the figure legends.

### Bone Marrow Reconstitution

Recipients, unless indicated otherwise were heterozygous CD45.2/CD45.1 to allow identification of donor control (CD45.1) and *Mrtfab*KO^T^ (CD45.2) cells. After watering on acidified water for >1 week, recipients were irradiated twice using ^137^Cs (6 Gy, 5 Gy for *Rag2*^-/-^ mice) with a 3hr interval; 10^6^ donor bone marrow cells were injected into the tail vein the following day, and animals analysed 6 weeks later. For negative selection experiments, cohorts of control or *RIP1-mOva* mice (CD45.2/CD45.1) were used.

### FACS analysis

Tissues were disaggregated through a 70μm nylon mesh in cold RPMI 10% FCS. Blood was collected into Sarstedt EDTA KE/1.3 tubes before red blood cell lysis. Cells were stained in ice-cold FACS buffer (1% FCS in PBS). 10μm latex beads (Coulter 6602796) were added to samples to allow normalisation for volume. DAPI staining was used to exclude dead cells in live cell analysis. For intracellular staining cells were stained with fixable viability dye (eBioscience 65-0865-14) and fixed using eBioscience Foxp3 staining buffer set (00-5523-00) for Foxp3 staining. Staining used combinations of fluorochrome-conjugated antibodies at a dilution of 1:200 unless stated otherwise. ***eBioscience*:** anti-CD4 (RM4-5), anti-CD8 (56-6.7), anti-CD11b (M1/70), anti-CD45.1 (A20), anti-CD45.2 (104), anti-CD49b (DX5), anti-CD117 (2B8), anti-CD122 (5H4), anti-CXCR4 (2B11), anti-Gr-1 (RB6-8C5), TCRβ (H57-597), anti-Qa2 (69H1.9-9), anti-Foxp3 (FJK-16s). ***Biolegend***: anti-CCR7 (4B12), anti-CCR9 (9B1), anti-H-2kb (AF6-88.5), anti-CD62L (MEL-14), anti-CD69 (H1.2F3), streptavidin Brilliant violet 421 (405225)***. BD Pharmingen***: anti-CD3ε (2C11), anti-CD24 (M1/69), anti-CD25 (7D4); anti-γδ TCR (GL3), anti-CD44 (IM7), anti-CD49b (DX5), anti-CD69 (H1.2F3), anti-B220 (RA3-6B2), anti-NK1.1 (PK136), anti-TCRβ (H57-597)***. R&D***: anti-S1P1 (FAB7089R).

### Cell purification and sorting

SP thymocytes or CD4 or CD8 T cells were purified by negative selection using MACS separation (Miltenyi Biotec 130104454, 130104075). Further purification of immature and mature thymocytes were identified by surface staining with anti-CD4, anti-TCRβ, anti-CD69 and anti-CD62L followed by cell sorting. T_regs_, γδ T cells, NK T cells and dead cells were excluded by staining with biotinylated anti-CD25, anti-γδ TCR and anti-DX5 and identified with streptavidin BV421 and DAPI. CD4+CD25-CD45RB^hi^ T_con_ and CD4+CD25+CD45Rb^lo^ T_reg_ cells were sorted from MACS-enriched CD4 populations. Sorts were performed on a BD INFLUX using Sortware software for analysis. Dendritic cells were prepared from Liberase TM (Roche) digested RAG2-/- spleens by CD11c+ MACS positive selection (Miltenyi Biotec 130108338).

### Homing assays and homeostatic proliferation

MACS purified T-cells from control and *Mrtfab*KO^T^ mice were labelled using 2μM CFSE (Molecular Probes) in RPMI, 37°C 10min or 1.5μM SNARF (Molecular Probes) in RPMI, 37°C, 20min, mixed 1:1, and 1×10^5^ cells injected into the tail vein of C57BL6 mice. Numbers of CFSE- and SNARF-positive cells present in the spleen and secondary lymph nodes were determined by FACS, and enumerated relative to input.

### BrdU labelling

Mice were injected intraperitoneally with 1mg BrdU, repeated after 4 hours. Tissues were harvested at the indicated time points after the second injection, cells surface stained to identify thymic populations and BrdU incorporation revealed using BD Pharmingen APC BrdU Flow kit (552598) according to manufacturer’s instructions.

### In vitro proliferation and survival assays

For CD3/CD28 stimulation, flat-bottom 96-well plates were coated overnight with the indicated concentration of anti-CD3 (2C11; in-house) and anti-CD28 (37.51; BD Biosciences) and washed with PBS and media. T cells (Total LN, FACS sorted or MACS-purified) were labelled with CFSE and plated, and the resulting CFSE dilution / proliferation profiles assessed by FACS.

For IL-7 and IL-15 survival and proliferation assays, CD2cre and *Mrtfab*KO^T^ LN CD8+ T cells were MACS purified by negative selection and labelled with CFSE. 10^5^ cells were plated in round-bottom 96-well plates in full medium supplemented with IL-7 or IL-15 (Peprotech 21714, 217-15), cultured for 5 days, and CFSE profiles analysed. Live cell recovery was determined using 10μm latex beads (Coulter 6602796).

### Thymic histology

Thymuses were fixed overnight in 10% buffered formalin followed by transfer into 70% Ethanol and embedded flat in paraffin. 3μm sections 250μm apart were stained with H&E for analysis of cortical and medullary staining. Images were acquired with a Zeiss AxioScan.Z1 slide scanner and quantitation was performed using QuPath software (Bankhead et al., 2017). Briefly, a pixel threshold was set to automatically detected the total tissue area and the medulla areas normalized to it for calculating the medullary area/over total tissue area. For analysis of the perivascular space, (Maeda et al., 2014) thymuses were frozen in OCT, sectioned and slides fixed in acetone before staining with anti-CD3 (2C11), anti-CD31 (EPR17259) and anti-ER-TR7 (Novus NB100-64932).

### Transwell migration

Transmigration assays were performed using permeable support polycarbonate membranes (Costar) 3μM or 5μM pore size. For S1P migration CD4SP thymocytes were purified by MACS negative selection. After resting for 45 minutes in RPMI, 1% BSA (fatty acid free), 1 x 10^5^ cells were placed in the upper chamber and allowed to migrate for the indicated times towards 100ng/ml, S1P (Sigma 2709) in the lower chamber (all in RPMI, 1% BSA). For chemokine induced migration, 1 x 10^6^ rested thymocytes or lymph node cells, or 1 x 10^5^ T-blasts were placed in the upper chamber previously coated overnight with 25μg/ml fibronectin (Merck F2006), and allowed to migrate for 120 min to: 100ng/ml SDF-1; 250 ng/ml (BIO-TECHNE 460-SD), CCL19 (BIO-TECHNE 440-M3), 250 ng/ml CCL21 (BIO-TECHNE 457-6C); in the lower chamber (all RMPI, 10% FCS unless otherwise indicated).

### F-actin content

Thymocytes (10^7^/ml) were rested for 45 minutes then stimulated with chemokines or S1P in RPMI 1% BSA, fixed for 30min using an equal volume of 4% PFA, then washed with FACS buffer and stained for cell surface markers and with FITC labelled phalloidin (1:50 (Invitrogen F432).

### Lacrimal gland and intestinal pathology

Tissue sections were fixed in 10% neutral buffered formalin (NBF) and embedded in paraffin. 3μm sections were stained with hematoxylin and eosin and anti-CD3 for histological interpretation. Infiltration of T cells and lymphocytes and into the lacrimal glands or large intestine was scored by Alejandro Suarez-Bonnet and Eleanor Herbert (Crick experimental histopathology facility) as follows: 0, not present; 1, mild; 2, moderate; 3 severe. Further staining of fixed lacrimal glands employed CD4 (abcam, ab183685, 1:750), CD8 (abcam, ab217344;1:500), Foxp3 (CST, 12653S;1:400), or B220 (BD Bioscience, 553086; 1:750). For fecal Lipocalin measurement, fecal pellets were collected and resuspended in 300ul 0.1% Triton PBS per tube. Solid material was removed by centrifugation and the supernatant assayed LCN2 using the R&D ELISA duo kit DY1857-05 according to manufacturers instructions

### Thymocyte transfer and in vivo suppression experiments

For thymocyte transfer assays 25 x 10^6^ Total thymocytes purified from pooled CD2-Cre or *Mrtfab*KO^T^ donors were injected intravenously into *Rag*2^-/-^recipients, with or without 2.5 x 10^6^ splenic derived sorted CD4+CD25+CD45RB^lo^ T_reg_ cells from wildtype animals, and analysed 10 weeks later. For *in vivo* suppression assays, 300,000 T_con_ and 150,000 T_reg_ cells were transferred intraperitoneally into *Rag1*^-/-^ or *Rag2*^-/-^ recipients. Animal weights were monitored twice weekly, and daily when disease was apparent. Tissues were harvested at defined times following transfer or upon cull of animals exhibiting persistent weight loss or failure to thrive. Formalin-fixed, paraffin-embedded sections were examined blind to experimental grouping by a board-certified Veterinary Pathologist, and histopathology semi-quantitatively assessed as: absence, 0; mild, 1; moderate, 2; severe, 3.

### *in vitro* suppression assays

For Ova peptide *in vitro* suppression assays, 50,000 CFSE labelled OT-II T_con_ cells from donor animals (3 mice pooled per genotype) were mixed in varying ratios with CD2cre and *Mrtfab*KO^T^ OT-II TCR transgenic T_reg_ in the presence of 5000 splenic DCs (prepared from RAG2^-/-^ splenocytes using CD11c microbeads, Miltenyl Biotec 130108338) and 0.1 µM OVA_323-339_ in round-bottom 96-well plates. CD25 upregulation and CD44 upregulation was analysed.

### qRT-PCR

Total thymocyte RNA was isolated using the GenElute kit (Sigma), DNA’ase I treated, and cDNA synthesised using the Superscript III Reverse Transcriptase kit (Life Technologies). cDNA was quantified using Sybr Green Express (Life Technologies) and an ABI 7900HT Fast RT-PCR instrument, normalizing to the abundance of RpS16 cDNA. Primers were:

Actb intronic: F, CGTAGCGTCTGGTTCCCAAT; R, GTGTGGGCATTTGATGAGCC

Actb exonic: F, CGCCACCAGTTCGCCAT; R, CTTTGCACATGCCGGAGC

Actg intronic: F, CTGGCCGAGGACATTTTCTG; R, GAAGAAGCCCCGGAATTAGC

Actg exonic: F, ATGGAAGAAGAAATCGCCGC; R, AGGGTCAGGATACCCCTCTT

Cxcr4 intronic: F, CTAGAACCGCTGTGTGTTGG; R, GCAGCGGACCCAGTTAGTTA

Cxcr4 exonic: F, AACCTCTACAGCAGCGTTCT; R, GATCCAGACGCCCACATAGA

Ccr7 intronic: F, AGCAAGAGTGGCCTTGATCT; R, GGGAACATCTAGCCTGGGTT

Ccr7 exonic: F, AGGCCTTGATCACCATCCAA; R, CACCTTGATGGCCTTGTTCC

S1pr1 intronic: F, ATGGGTGGCTCTCATTTCCA; R, AAACCTTCCCTTGGGTCACT

S1pr1 exonic: F, TGAACTCAGGTACCAACCCC; R, TTGGGGCATTTGCAACAAGA

RpS16: F, CGCACGCTGCAGTACAAGTTACT; R, ACATGTCCACCACCCTTCACAC

### Gene deletion validation

DNA was prepared using QuickExtraction DNA solution (LGC Biosearch Technologies) and analsed by PCR, using the following genotyping primers.

***iCre***: F, TCAACATGCTGCACAGGAGAT; R, ACCATAGATCAGGCGGTGGGT

***Mrtfa***: GTTGCTCAGTCATGTGACACCTGTACAG (GT5), with either GT6: GGCTTCAGTACCTTCCTAAGCTCTGCAG (GT6; wildtype *Mrtfa* allele, 522bp) or ATGGTGGATCCTGAGACTGGCGAATTC (LacZ3-QB; knockout *Mrtfa* allele; ∼350bp)

***Mrtfb***: CATGGCGACTTCCTTCTCCTCTTCTCAAGGCTG (SAF), with GGCTTAGACAAGATGGTTGGTCTGGCACTGC (EXR; WT *Mrtfb* allele, 460bp; floxed *Mrtfb* allele 570bp); or with CCAGTGGTGTCCAGTCTTACTGAACAGCTCACTCAG (LAR; *Mrtfb* deleted allele, 364bp).

MRTF expression was monitored using anti-MRTF-A (in-house) anti-MRTF-B (in house) with anti-pan ERK (BD bioscience 610124) control.

## SUPPLEMENTARY FIGURE LEGENDS

**Figure S1.**
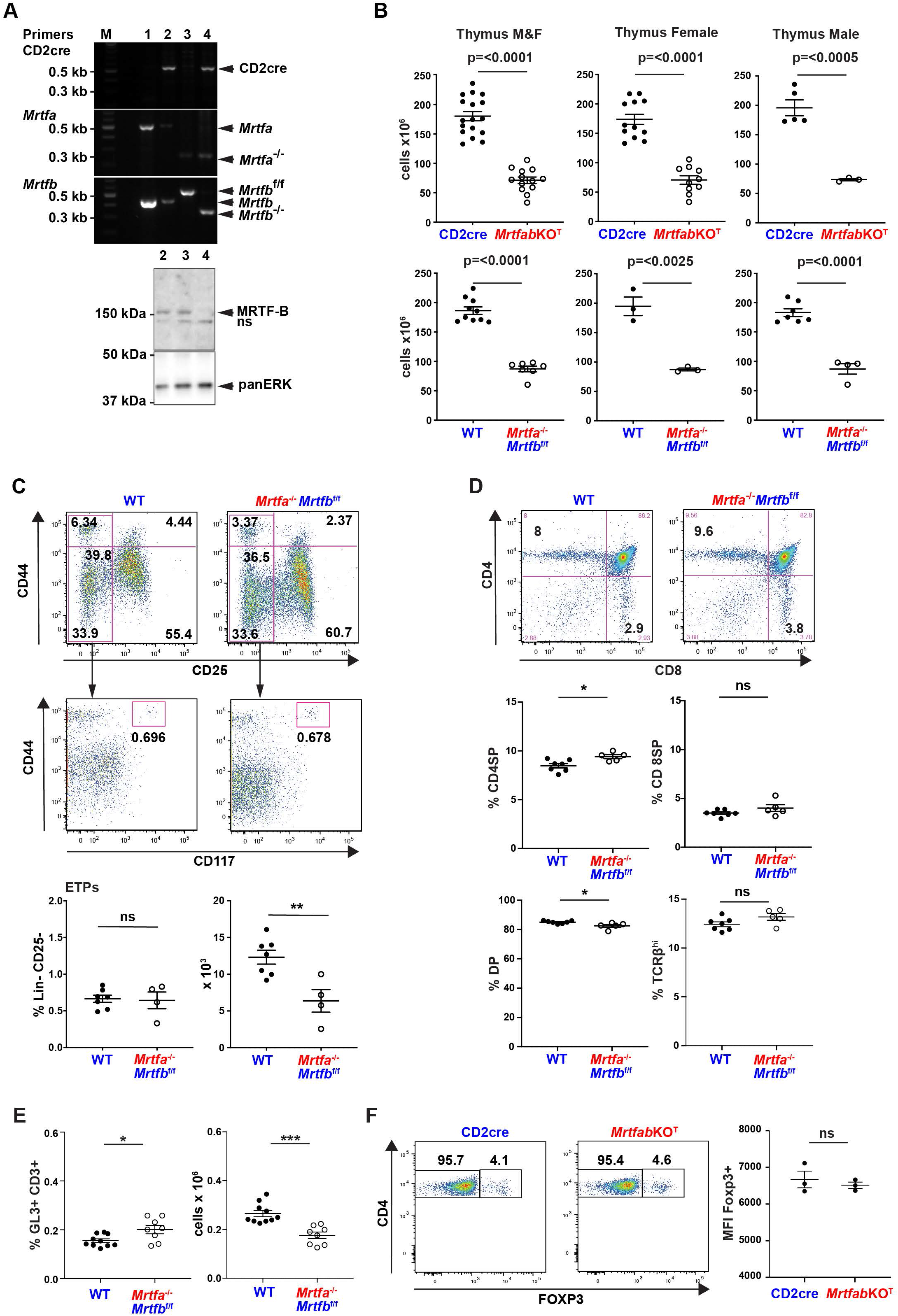
Normal thymocyte positive selection in *Mrtfab*KO^T^ animals. A. Top, genomic PCR analyis of DNA extracts from young adult (1) WT, (2) *CD2Cre*, (3) *Mrtfa*^-/-^*Mrtfb^f/f^*, and (4) *Mrtfab*KO^T^ thymocytes. Bottom, immunoblot analysis of thymocyte extracts. B. Thymocyte cellularity of age-matched CD2cre and *Mrtfa*^-/-^ *Mrtfb^f/f^* mice. Left, all animals; right, male and female animals. Thymic cellularity of age-and sex-matched populations was as follows: all CD2cre, 180.2 ± 32.6 (n=17); all *Mrtfab*KO^T^, 71.32 ± 19.9 (n=13); female CD2cre, 173.6 ± 32.42 x 10^6^ (n=12); female *Mrtfab*KO^T^, 70.64 ± 3.2 x 10^6^ (n=10); male CD2cre, 196.0 ± 30.8 x 10^6^ (n=5); male *Mrtfab*KO^T^, 73.6 ± 3.2 x 10^6^ (n=3); all WT, 186.3 ± 20 (n=10); all *Mrtfa*^-/-^ *Mrtfb^f/f^*, 87.19 ± 12.9 (n=7); female WT, 194.7 ± 27.3 x 10^6^ (n=3); female *Mrtfa*^-/-^ *Mrtfb^f/f^,* 87.3 ± 4.2 x 10^6^ (n=3); male WT, 182.7 ± 17.4 x 10^6^ (n=7); male *Mrtfa*^-/-^ *Mrtfb^f/f^*, 87.1 ± 17.92 x 10^6^ (n=4). C. ETPs in age-matched *Mrtfa*^-/-^*Mrtfb^f/f^* male mice (from Figure S1B). Top, representative live-gated, lineage negative CD44 CD25 profiles; middle. CD17 CD44 profiles of the CD25-CD44^lo^ populations, with CD44+ CD117+ ETPs indicated; bottom, ETP quantitation (data combined from two independent experiments, at least two mice each genotype). D. Analysis of thymic populations of age and sex matched WT and *Mrtfa*^-/-^*Mrtfb^f/f^* mice. Top, representative live-gated CD4 and CD8 profiles of WT and *Mrtfa*^-/-^*Mrtfb^f/f^* thymocytes; middle, CD4 and CD8 expression; bottom, DP and TCRβ expression (data combined from two independent experiments, at least two mice each genotype). E. Thymic GL3+ CD3+ γδ T cells. Proportion and numbers of thymic GL3+ CD3+ γδ T cells in *Mrtfa*^-/-^ *Mrtfb^f/f^* mice (data combined from three independent experiments, at least two mice each genotype (WT n=10, *Mrtfab*KO^T^ n=8)). F. Left representative live-gated CD4 FoxP3 profiles of CD2cre and *Mrtfab*KO^T^ thymocytes. Right, relative FoxP3 MFI of CD2cre and *Mrtfab*KO^T^ CD4+thymocytes. Data combined from three 3 mice each genotype, analysed in single experiment.

**Figure S2.**
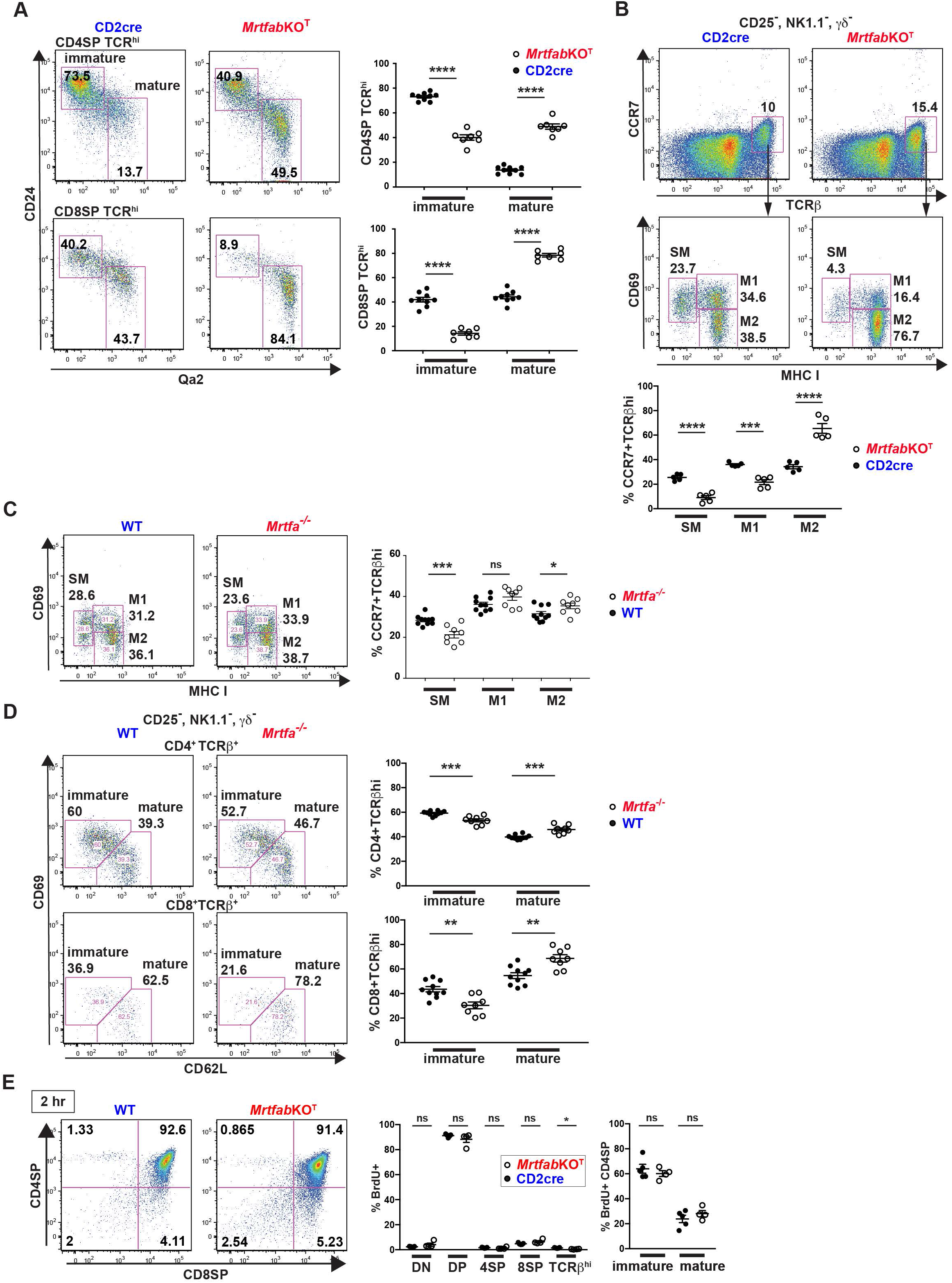
Egress of mature thymocytes is impaired in *Mrtfab*KO^T^ animals. A. Representative CD24 Qa2 profiles of live-gated CD4SP TCR^hi^ and CD8SP TCR^hi^ thymocytes in CD2cre and *Mrtfab*KO^T^ mice. Immature (CD24^hi^ Qa2-) and mature (CD24^lo^ Qa2+) populations are quantified at the right. Data are combined from two independent experiments, at least two mice each genotype (CD2cre, n=8; *Mrtfab*KO^T^, n=5). B. Representative CCR7 TCRβ profiles of live-gated, CD25-, GL3-, NK1.1-thymocytes in CD2cre and *Mrtfab*KO^T^ animals, with the CCR7+, TCRβ^hi^ populations analysed for CD69 and MHC I below. Thymocytes defined as semi-mature (SM), CD69^hi^ MHC I-; mature 1 (M1), CD69^hi^ MHC I+; mature 2 (M2), CD69^lo^ MHC I+. CCR7+ TCRβ^hi^ populations, are quantified below. Data are combined from three independent experiments, 1-2 mice of each genotype (CD2cre, n=5; *Mrtfab*KO^T^, n=5). C. Maturing thymic populations of age-matched WT and *Mrtfa^-/-^Mrtfb^fl/fl^*mice, defining SM, M1 and M2 populations using CD69 and MHC I profiles of live-gated, CD25-, GL3-NK1.1-CD4+ TCRβ^hi^ WT and *Mrtfa^-/-^Mrtfb^fl/fl^* thymocytes as in (B). Left, representative CD69, MHC I profiles. Right, data are combined from three independent experiments, at least two mice each genotype (WT, n=10; *Mrtfa^-/-^*, n=7). D. Maturing thymic populations of age-matched WT and *Mrtfa^-/-^Mrtfb^fl/fl^*mice, defining immature and mature populations using CD69 and CD62L profiles of the animals analysed in (C). Left, representative CD69, CD62L profiles. Right, proportions of immature and mature cells. E. WT and *Mrtfab*KO^T^ animals given two injections of BrdU, and thymocytes analysed 2h after the second injection. Left, representative BrdU+ gated CD4 and CD8 profiles. Centre, proportion of BrdU+ cells in DN, DP, CD4SP, CD8SP and TCRβhi populations. Right, proportions of BrdU+ thymocytes in immature CD69^hi^ CD62^lo^ and mature CD69^lo^ CD62L^hi^ TCRβ^hi^ CD4SP populations. Data from 5 CD2cre and 4 *Mrtfab*KO^T^ mice.

**Figure S3.**
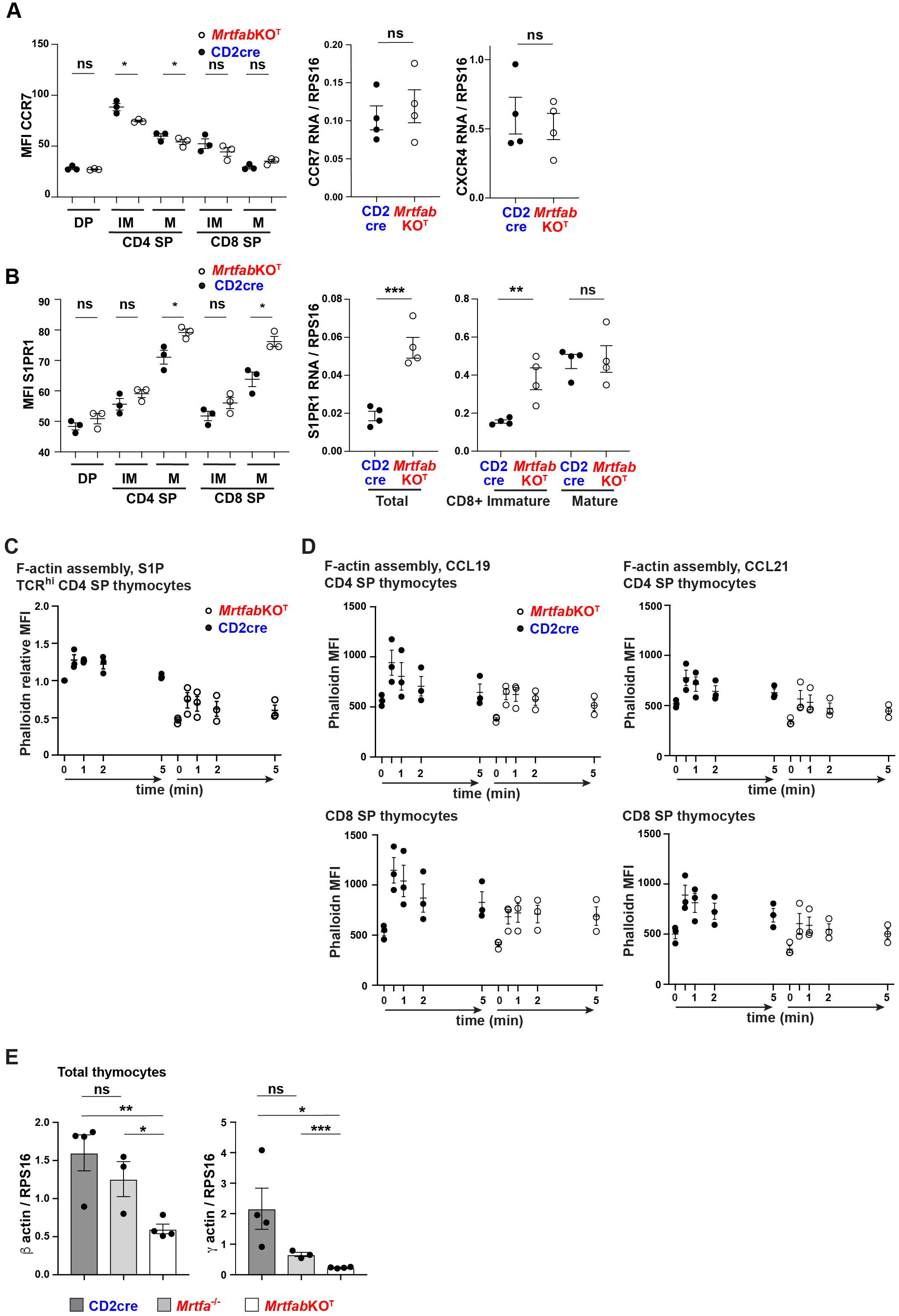
Migration defects in *Mrtfab*KO^T^ thymocytes. A. CCR7 and CXCR4 expression on thymocyte subsets. Left, live-gated thymocytes from CD2cre and *Mrtfab*KO^T^ mice were stained for CD4 and CD8, maturity markers CD62L, CD69 and CCR7. Data are of three age matched male mice of each genotype. Right, thymocyte RNA was analysed by qRT-PCR for CXCR4 or CCR7 and quantified relative to RPS16. Data are mean ± SEM (n=4 female mice of each each genotype). B. S1PR1 expression on thymocyte subsets. Left, live-gated thymocytes from CD2cre and *Mrtfab*KO^T^ mice were stained for CD4 and CD8, S1PR1, and maturity markers CD62L and CD69. Data are of three age-matched male mice of each genotype. Right, thymocyte RNA was isolated and analysed by qRT-PCR as in (A) for S1PR1 and quantified relative to RPS16. C. CD4 SP-enriched thymocytes from CD2cre and *Mrtfab*KO^T^ animals were stimulated with S1P and stained for TCRβ and F-actin. F-actin levels are normalised to the unstimulated F-actin baseline in CD2cre thymocytes. Data are mean ± SEM from three independent experiments. D. Thymocytes were pooled from 3 CD2cre and 2 *Mrtfab*KO^T^ animals, stimulated with CCL19 or CCL21, and stained for CD4, CD8 and F-actin. Data are technical replicates ± SEM. E. Total thymocytes were isolated from CD2cre, *Mrtfa^-/-^Mrtfb^fl/fl^* and *Mrtfab*KO^T^ mice and β and γ actin mRNA levels analysed by qRT-PCR, expressed relative to RPS16. Data are mean ± SEM (CXD2cre, n=4; *Mrtfa^-/-^Mrtfb^fl/fl^*, n=3; *Mrtfab*KO ^T^, n=4).

**Figure S4.**
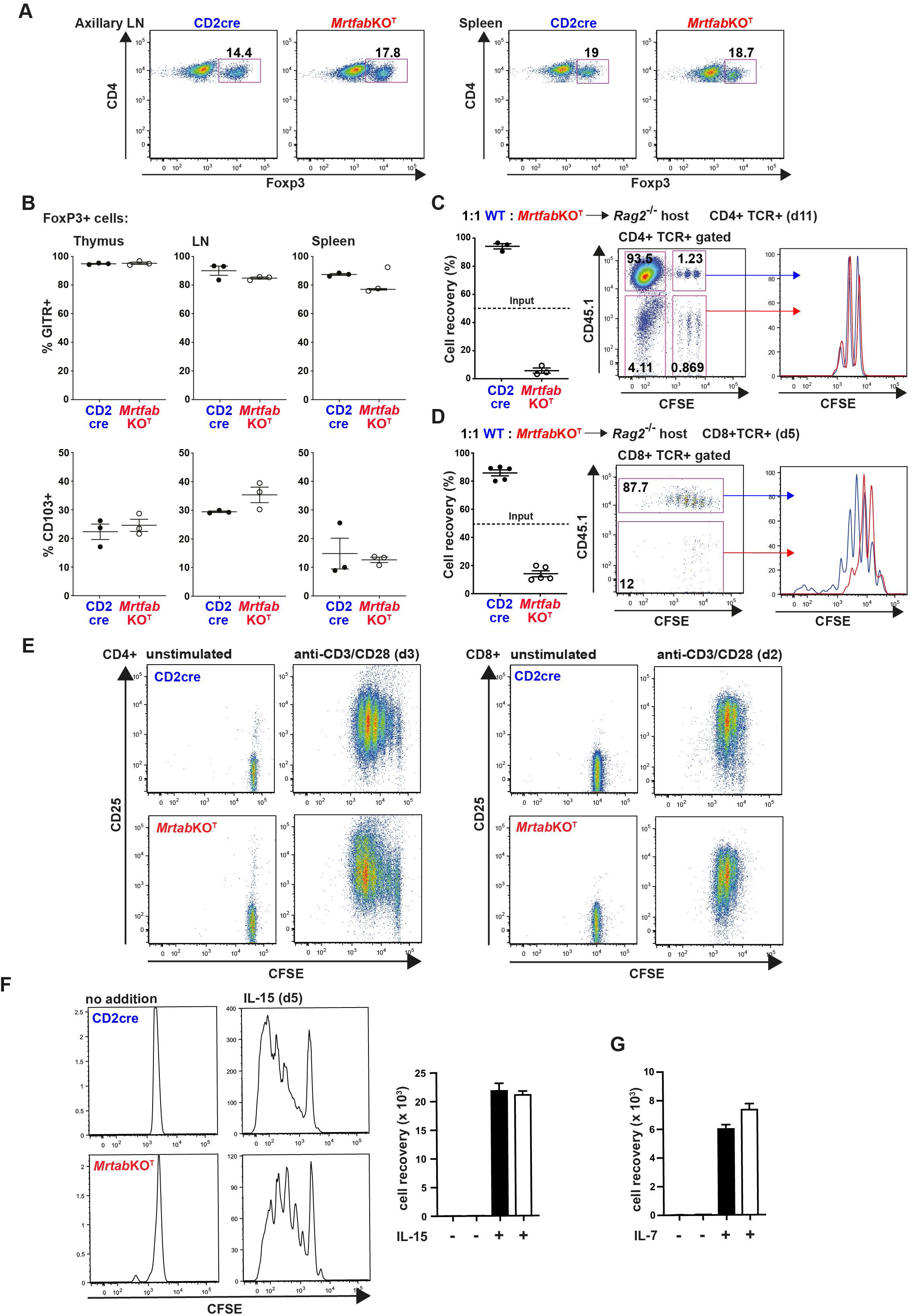
Lymphopenia in *Mrtfab*KO^T^ animals. A. Representative live-gated CD4 FoxP3 profiles of CD2cre and *Mrtfab*KO^T^ thymocytes in axillary lymph nodes and spleen. Data combined from three 3 mice each genotype, analysed in single experiment. B. Proportion of CD2cre or *Mrtfab*KO^T^ FoxP3+ T_reg_ cells expressing GITR or CD103 (integrin αE) in lymphoid organs. Data mean ± SEM, n=3. C. WT CD45.1 and *Mrtfab*KO^T^ CD45.2 T cells from 4 mice of each genotype were pooled, labelled with CFSE, co-transferred into Rag2^-/-^hosts at 1:1 ratio, and analysed 11 days later. Left, relative proportions of WT and *Mrtfab*KO^T^ CD4 cells recovered from pooled lymph nodes. Data are mean ± SEM, n=3 mice. Right, total CD4+ TCR+ gated CFSE CD45.1 staining, and comparison of the CD45.1+ (WT) and CD45.1-(*Mrtfab*KO^T^). A representative analysis is shown. D. WT CD45.1 and *Mrtfab*KO^T^ CD45.2 T cells were co-transferred into Rag2^-/-^ hosts at 1:1 ratio as in (B), and analysed 5 days later. Left, relative proportions of WT and *Mrtfab*KO^T^ CD8 cells recovered from pooled lymph nodes. Data are mean ± SEM, n=5 recipients, combined from two independent experiments, in one of which cells were labelled with CFSE before transfer (two recipients). Right, total CD8+ TCR+ gated CFSE CD45.1 staining, and comparison of the CD45.1+ (WT) and CD45.1-(*Mrtfab*KO^T^). A representative analysis of two recipients is shown. E. CFSE-labelled CD2cre and *Mrtfab*KO^T^ T cells were cultured on plate-bound αCD3/αCD28. Data are CFSE CD25 profiles of unstimulated cells and cells stimulated for 72hrs (CD4+) or 48hrs (CD8+). A representative of two independent experiments is shown. F. CFSE-labelled CD2cre and *Mrtfab*KO^T^ CD8+ T cells were cultured for 5 days with or without 100ng/ml rIL-15. Right, cell recovery. Data are mean ± SEM of three technical replicates. G. Recovery of CD2cre and *Mrtfab*KO^T^ CD8+ T cells after 5 days culture with or without 100ng/ml rIL-7. Data are mean ± SEM of three technical replicates.

**Figure S5.**
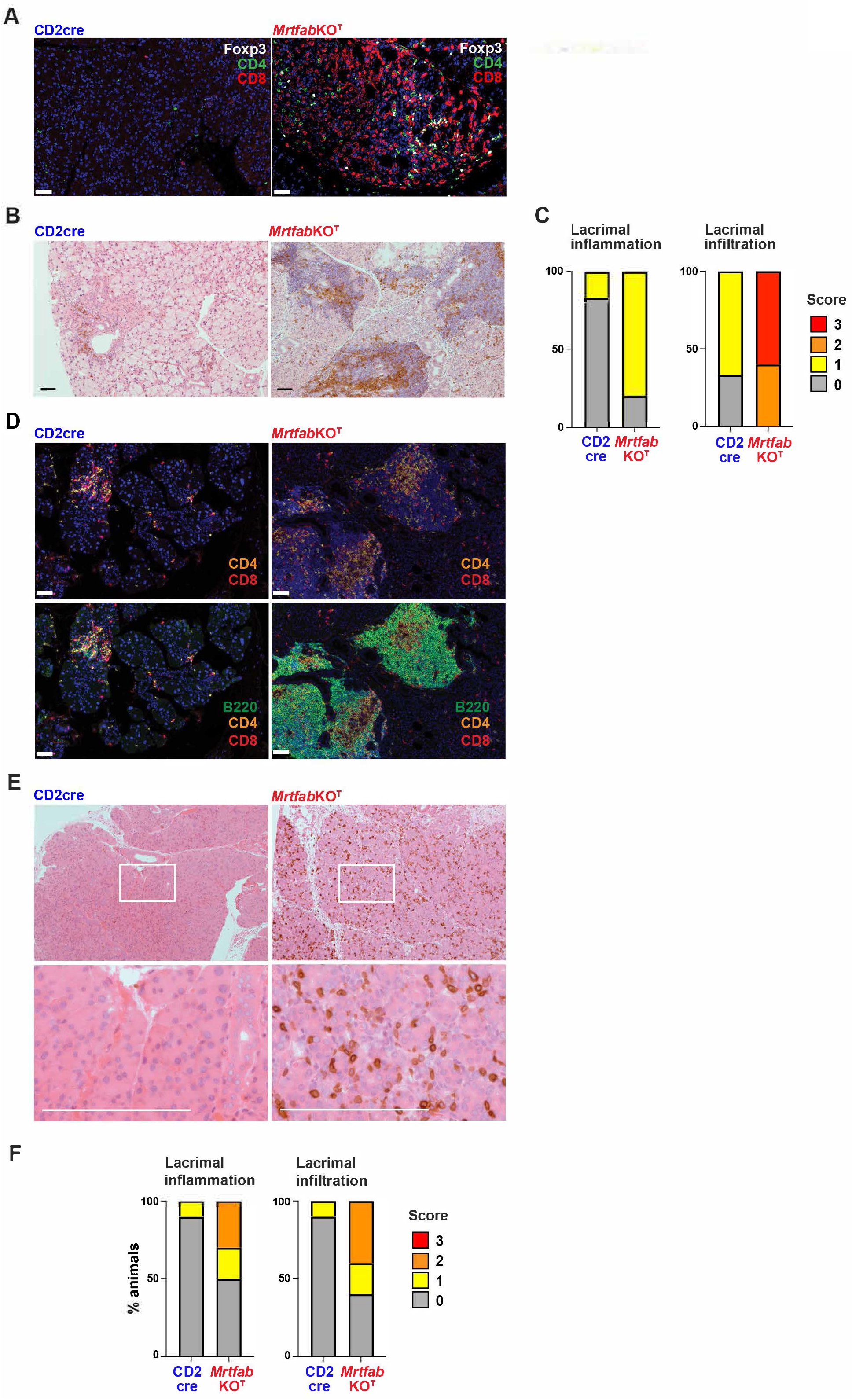
Autoimmune pathology in *Mrtfab*KO^T^ mice. A. Lacrimal gland histology from 4 week old male CD2Cre and *Mrtfab*KO^T^ mice stained for FoxP3, CD4 and CD8. Scale bar, 50μm. 3 out of 4 *Mrtfab*KO^T^ lacrimal glands displayed lymphocyte infiltration, with no infiltration observed in CD2cre lacrimal glands. B. 3μm lacrimal gland sections from older CD2cre and *Mrtfab*KO^T^ female mice (median ages 35 and 31 weeks respectively), were stained with (top) hematoxylin and eosin, and anti-CD3 (brown); scale bar, 50μm. *Mrtfab*KO^T^ pathology was graded as mild for inflammation and severe for infiltration in (C). C. Qualitative grading of lacrimal gland inflammation and lacrimal gland infiltration in older female mice as in B. CD2Cre (n = 6) and *Mrtfab*KO^T^ (n = 5) mice. D. 3μm lacrimal gland sections from 35 week old CD2Cre and 31 week old *Mrtfab*KO^T^ female mice were stained for CD4 and CD8 to detect T cell infiltrates, and B220 to detect extrafollicular B cells. Scale bar, 50μm. ^E.^ Lacrimal gland pathology is T cell dependent. Total thymocytes from CD2cre or *Mrtfab*KO^T^ mice (2 donors each genotype) were transferred into *Rag2*^-/-^ recipients (5 recipients per donor, 25 x 10^6^ cells each recipient) and pathology analysed 10 weeks after transfer. Images are lacrimal gland sections stained with H&E and anti-CD3 (brown). Image is from *Mrtfab*KO^T^ animal was graded moderate for lacrimal gland inflammation and infiltration in (F). Boxed sections of images are enlarged below. Scale bar, 1mm. F. Qualitative grading of lacrimal gland inflammation and infiltration from the experiment shown in (E).

**Figure S6.**
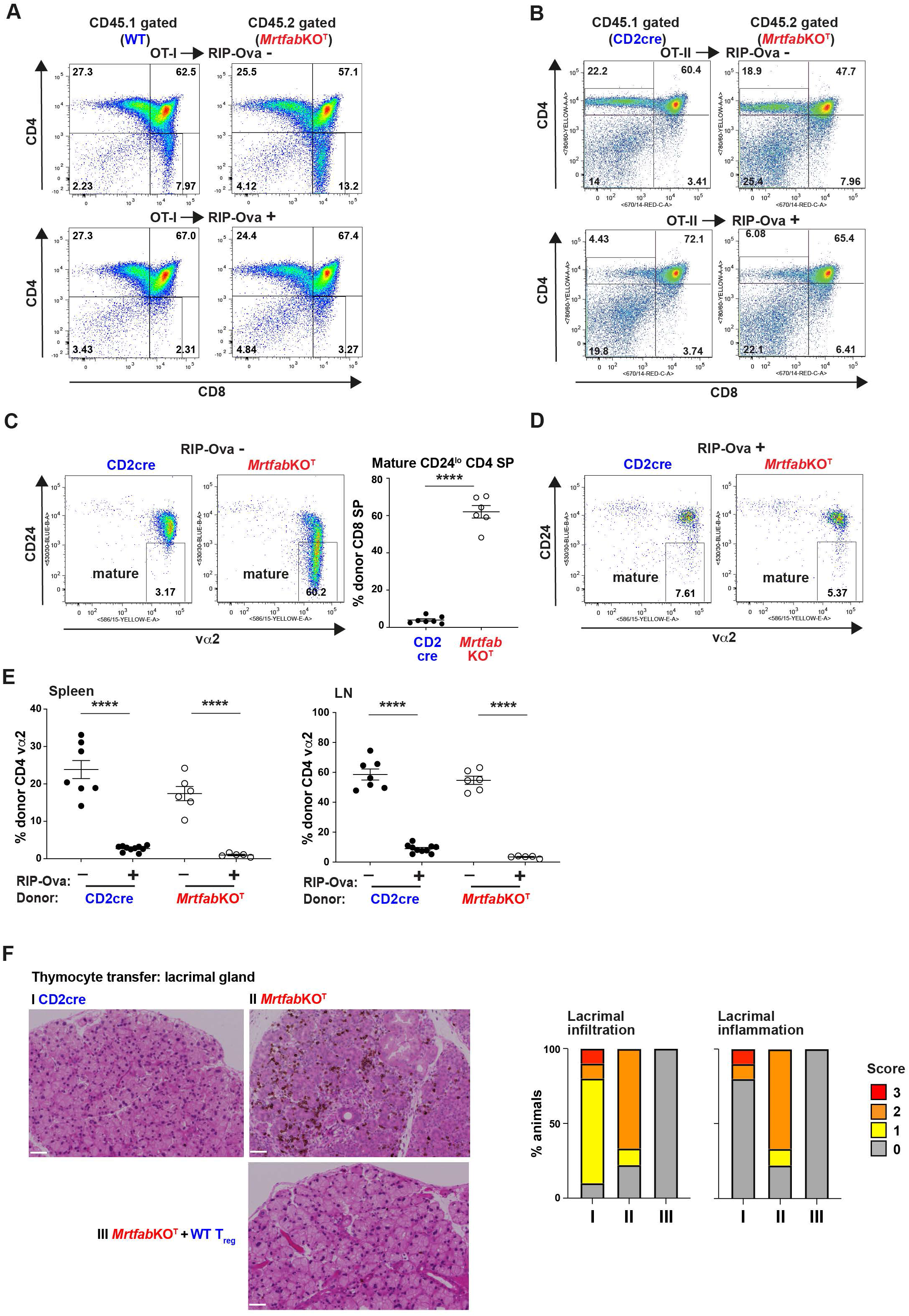
Negative selection is intact in *Mrtfab*KO^T^ mice. A. Irradiated female wildtype CD45.1/CD45.2 hosts were reconstituted with OT-I TCR transgenic bone marrow from either wildtype (CD45.1) or *Mrtfab*KO^T^ (CD45.1) animals. Data are representative CD4 / CD8 profiles of thymocytes from OT-I CD2cre and OT-I *Mrtfab*KO^T^ animals, gated on CD45.1 and CD45.2 respectively. Upper panels, wildtype host background, (Vα2 CD24 profile shown in Figure 6A); lower panels, RIP-mOva+ transgenic host background, Vα2 CD24 profile shown in Figure 6B. B. Irradiated female wildtype CD45.1/CD45.2 hosts were reconstituted with OT-II TCR transgenic bone marrow from either CD2cre (CD45.1) or *Mrtfab*KO^T^ (CD45.1) animals. Data are representative CD4 / CD8 profiles of thymocytes from OT-II CD2cre and OT-II *Mrtfab*KO^T^ animals, gated on CD45.1 and CD45.2 respectively. Upper panels, wildtype host background (Vα2 CD24 profile shown in Figure 6D); lower panels, RIP-mOva transgenic host background (Vα2 CD24 profile shown in Figure 6E). C. Left, representative CD24 / vα2 profiles of T cells from primary young adult CD2cre OT-II and *Mrtfab*KO^T^ OT-II transgenic animals. Right, proportions of mature CD4SP CD24^lo^ thymocytes in CD2cre and *Mrtfab*KO^T^ animals. Data are combined from three independent experiments (CD2cre, n=7; *Mrtfab*KO^T^, n=6). D. Representative CD24 / vα2 profiles of T cells from primary CD2cre (n=10) and *Mrtfab*KO^T^ (n=5) OT-II TCR RIP-mOva+ transgenic animals. E. Proportions of donor gated CD4+ vα2+ T cells in spleen and lymph nodes from the experiments in (C) and (D). F. T_reg_ cells suppress lacrimal gland pathology arising in *Rag2*^-/-^ recipients after total *Mrtfab*KO^T^ thymocyte transfer. Images are of 3μm lacrimal gland sections stained with hematoxylin and eosin, and anti-CD3 (brown), made 10 weeks after transfer of 25 x 10^6^ CD2cre or *Mrtfab*KO^T^ total thymocytes, with or without co-transfer of 0.25 x 10^6^ WT T_reg_ cells (CD4+, CD25+ CD45RB ^lo^). *Mrtfab*KO^T^ image is from an animal graded moderate for lacrimal gland inflammation and infiltration. Scale bar, 50μm. Right, qualitative grading of lacrimal gland inflammation and infiltration. CD2cre, n = 10; *Mrtfab*KO^T^, n = 10, *Mrtfab*KO^T^ + WT T_reg_, n=10.

**Figure S7.**
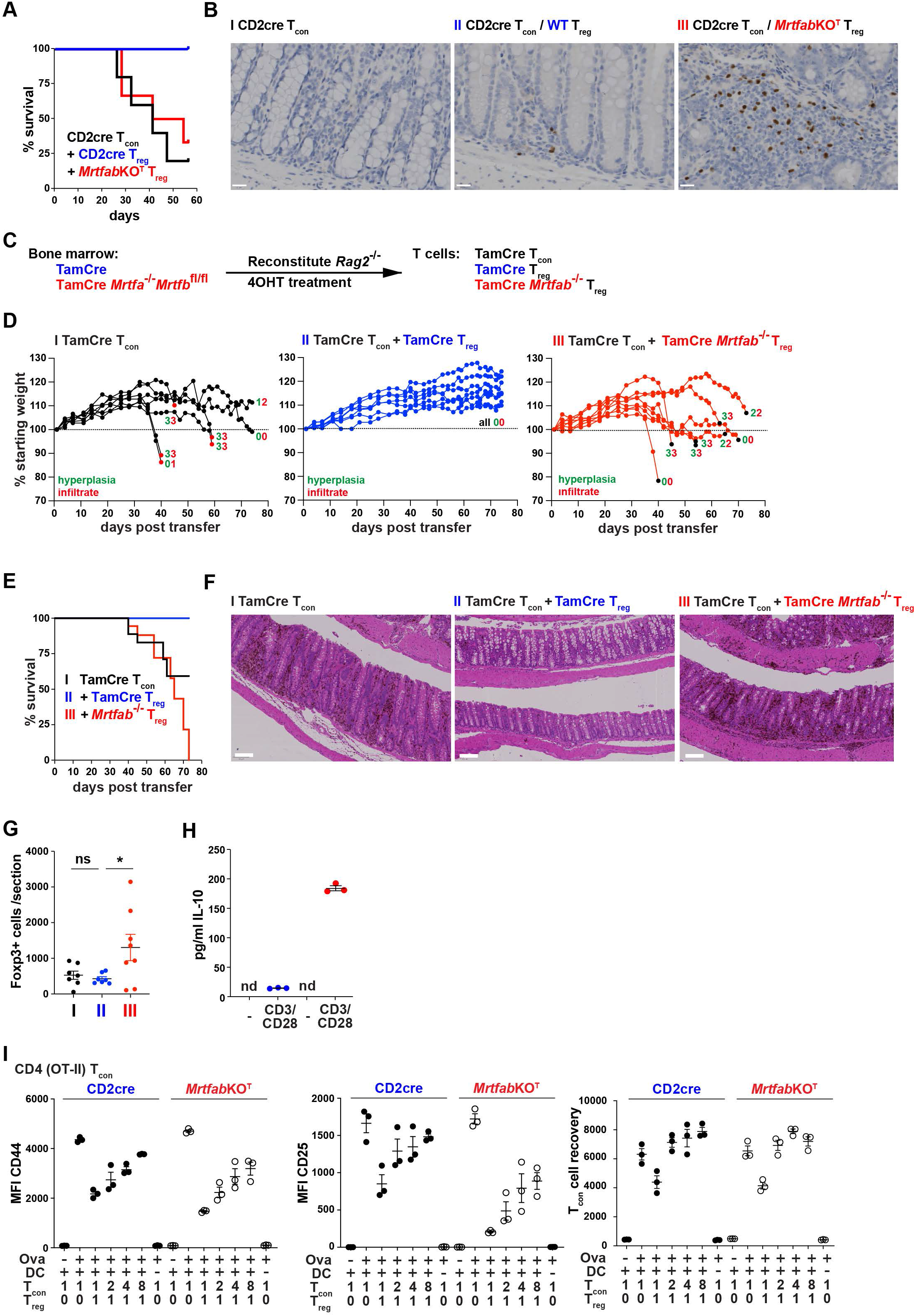
Autoimmune pathology in *Mrtfab*KO^T^ mice. A. Mortality following transfer of 300,000 CD2cre T_con_ (CD4+ CD25-CD45RB^hi^) cells, either alone or with 150,000 CD2cre or MrtfabKO^T^ T_reg_ (CD4+ CD25+ CD45RB^lo^) cells into male Rag2^-/-^ recipients. Animals were culled upon loss of >15% starting weight or persistent loss of condition. For individual weights and pathology see Figure 7A-D. B. FoxP3+ cells in the large intestine at following T_con_/T_reg_ transfer as in (A). 10x images of 3μm large intestine sections were made 10 weeks after transfer and stained with hematoxylin and anti-FoxP3 (brown). Scale bars, 20μm. C. Schematic outline of protocol for peripheral acute inactivation of the MRTFs and preparation of T_con_ and T_reg_ cells. D. Male *Rag1*^-/-^ recipients were injected with 300,000 TamCre T_con_ (CD4+ CD25-CD45RB^hi^) cells either alone, or with 150,000 TamCre or TamCre *Mrtfab*^-/-^ T_reg_ (CD4+ CD25+ CD45RB^lo^) cells, generated by peripheral inactivation of *Mrtfb* in *Mrtfa*^-/-^ *Mrtfb*^fl/fl^ cells using Cre^ERT^ (Maurice et al., 2024). Animals were culled upon loss of >15% starting weight or persistent loss of condition. For pathology summary, see Figure 7F. E. Individual animal weights and mortality in the experiment shown in (D). F. Mortality in the experiment shown in (D) G. Intestinal Foxp3+ cell numbers in the experiment shown in (D). H. Sorted CD4+ CD25+CD45RB^lo^ T_reg_ cells were cultured for 18hrs on plate bound anti-CD3 and anti-CD28, and IL-10 present in supernatants assayed by ELISA. Data are representative of two independent experiments; nd, not detectable. I. Activation of OT-II CD4+ conventional T cells by T_reg_ cells. T_con_ and T_reg_ cells were purified from CD2cre or *Mrtfab*KO^T^ OT-II TCR transgenic mice and mixed as indicated with *Rag2*^-/-^ splenic DCs and 0.1 µM mOva peptide. Upregulation of CD44 and CD25 and total T_con_ cell numbers were analysed 72h later. Data are mean ± SEM, from three replicate determinations.

